# A Transcriptomic Atlas of Healthy Human Skin Links Regional Identity to Inflammatory Disease

**DOI:** 10.1101/2025.05.10.653184

**Authors:** Sahiti Marella, Rachael Bogle, Jennifer Fox, Lam C. Tsoi, Xianying Xing, Yiqian Gu, Joseph Kirma, Mrinal K. Sarkar, Vincent van Drongelen, Benjamin Klein, Jeff H. Kozlow, Paul W. Harms, Katherine Gallagher, Shruti Naik, Vito W. Rebecca, Bogi Andersen, Mio Nakamura, J. Michelle Kahlenberg, Robert L. Modlin, Allison C. Billi, Johann E. Gudjonsson

## Abstract

Human skin is not a uniform organ but a mosaic of anatomically distinct niches, with each site finely tuned to unique environmental demands and immune pressures. Yet, the molecular determinants that define these regional identities and their relationship to site-specific vulnerability to inflammatory disease remain poorly understood. Here, we generate a high-resolution single-cell atlas of human skin, profiling 274,834 cells from 96 healthy samples across 7 anatomically distinct sites (acral, arm, axilla, back, face, leg and scalp). Our analysis reveals striking region-specific transcriptional and cellular networks, uncovering how local immune-stromal crosstalk governs tissue homeostasis and underpins anatomical susceptibility to distinct inflammatory diseases such as such as systemic lupus erythematosus (SLE), atopic dermatitis (AD), and psoriasis. These findings illuminate the tissue-intrinsic foundations of regional immune identity and provide a blueprint/resource for the development of precision therapies tailored to the distinct immunological microenvironments of specific anatomical skin sites.

## INTRODUCTION

A striking and poorly understood feature of many inflammatory skin diseases, such as systemic lupus erythematosus (SLE), psoriasis, and atopic dermatitis (AD), is their predilection for specific anatomical sites. SLE affects an estimated 204,000 individuals in the United States alone, and psoriasis and AD impact 1–3% and 2.6% of the global population respectively^1–4^. While cutaneous manifestations of SLE disproportionately affect the face and scalp, psoriasis often manifests on the face, scalp, and acral (palm and sole) regions, and AD typically occurs on the arm, face, leg, and scalp^5–11^. Uncovering the molecular and cellular networks that drive the site-specificity of these diseases could enable the development of precision therapeutic strategies tailored to the distinct immune-stromal microenvironments in different anatomical sites.

The skin is a structurally complex organ, composed of the epidermis, dermis, and hypodermis layers. Together, cells within these layers exert specialized functions to maintain the protective barrier between the body and the external environment, provide immune surveillance, and promote wound healing, sensory changes, and thermoregulation^12,13^. The epidermis, the outermost layer of the skin, primarily contains keratinocyte (KC) subtypes, pigment-producing melanocytes, and antigen-presenting Langerhans cells^13^. KCs are the predominant cell type of the epidermis and produce key structural proteins, including keratins, to form the primary physical barrier of the skin^14^. Directly beneath it is the dermis, composed of eccrine and sweat glands, hair follicle cells, fibroblasts, endothelial cells, pericytes and Schwann cells^13^. Dermal fibroblasts produce important components of the extracellular matrix (ECM), such as collagens, elastic fibers, and basement membrane associated macromolecules, which together act as scaffolds and are essential for maintaining skin structure^13,15^. The deepest skin layer, the hypodermis, includes adipose lobules, additional sensory neurons, and blood vessels^13^. Skin-resident immune cells, such as Langerhans cells, macrophages, T cells, in the epidermis and dermis, help maintain immune surveillance and regulate inflammation^13^.

Anatomical site-dependent specialization of skin structure and function arises in response to stimuli such as varying exposure to UV radiation, moisture levels, mechanical stress, friction, and microbial colonization^16,17^. These stimuli require skin to undergo adaptations, which include changes in structural organization, cellular composition, and gene expression signatures across different anatomical sites^16,18^. For example, acral skin must withstand increased mechanical stress and demonstrates unique transcriptional signatures associated with stress-response pathways, while facial skin is subjected to recurrent UV exposure and shows enhanced regulation of immune and inflammatory pathways^19,20^. These site-specific differences likely have a role in the striking regional specificity observed in inflammatory skin diseases such as SLE, psoriasis, and AD. Several transcriptomics studies have furthered our understanding of the key cellular players and molecular alterations that drive these diseases^21–23^. However, a comparative atlas of healthy skin across disease-relevant anatomical regions, necessary for uncovering the mechanisms which govern regional disease susceptibility, remains lacking. To address this gap, we integrated single-cell and spatial transcriptomics to profile healthy skin across anatomically distinct sites including acral, arm, axilla, back, face, leg and scalp. This atlas provides foundational insights into the cellular and molecular landscape across anatomical sites, offering critical context for understanding regional disease susceptibility and serving as an important resource for future investigations into skin homeostasis and disease pathogenesis, and to guide precision dermatology.

## RESULTS

### A single-cell atlas of healthy human skin reveals anatomical site-specific differences in cellular composition

To establish a comprehensive reference of anatomical site-specific skin cellular diversity, we performed scRNA-seq on skin samples from 96 healthy samples spanning seven anatomical sites including acral, arm, axilla, back, face, leg and scalp (Fig. 1a). The resulting quality-controlled healthy skin atlas contained a total of 274,834 cells (Fig. 1b) with an average of 2432 genes and 7939 transcripts detected per cell. The donor cohort included individuals with a balanced male-to-female ratio (Fig. 1c) and across a wide age range (19-93 years; Fig. 1d), ensuring broad representation of skin diversity in the general population (Supplementary Table S1). To identify distinct cellular populations, we applied unsupervised clustering followed by Uniform Manifold Approximation and Projection (UMAP), which revealed all major skin cell types, including Keratinocytes (KCs), Follicle Cells, Fibroblasts, Pericytes, Melanocytes, Endothelial Cells, L Endothelial Cells, Eccrine Cells, Schwann Cells, Mast Cells, T Cells, and Myeloid Cells (Fig. 1e). We validated our cluster annotations by comparing top cluster-specific differentially expressed genes with the canonical cell type-specific genes reported in previous scRNA-seq studies (Supplementary Fig. 1a; Supplementary Table S2)^22,24–27^. KCs and fibroblasts were the most abundant cell populations across all anatomical sites, with T cell and myeloid cell populations exhibiting regional differences in abundance (Fig. 1f). Notably, the relative cell type proportions in facial skin exhibited the greatest divergence from other anatomical sites (Fig. 1f). To further contextualize our scRNA-seq findings, we incorporated CosMx spatial transcriptomics, to visualize the spatial organization and proximity of major skin cell types across anatomical sites (Fig. 1g-m). These spatial maps confirmed the expected localization of KCs in the epidermis, fibroblasts and immune cells in the dermis, and melanocytes along the basal epidermal layer. Notably, acral skin displayed a thickened epidermis with abundant keratinocyte populations, while facial and scalp skin showed enriched immune cell infiltration and stromal complexity, supporting the transcriptional distinctions we observed in scRNA-seq (Fig. 1g-m). When comparing transcriptional profiles across anatomical sites, we identified genes consistently expressed across multiple sites as well as distinct gene expression signatures unique to specific bodysites, highlighting both shared and regionally specialized transcriptional programs (Supplementary Table S3a-c).

**Figure 1.**
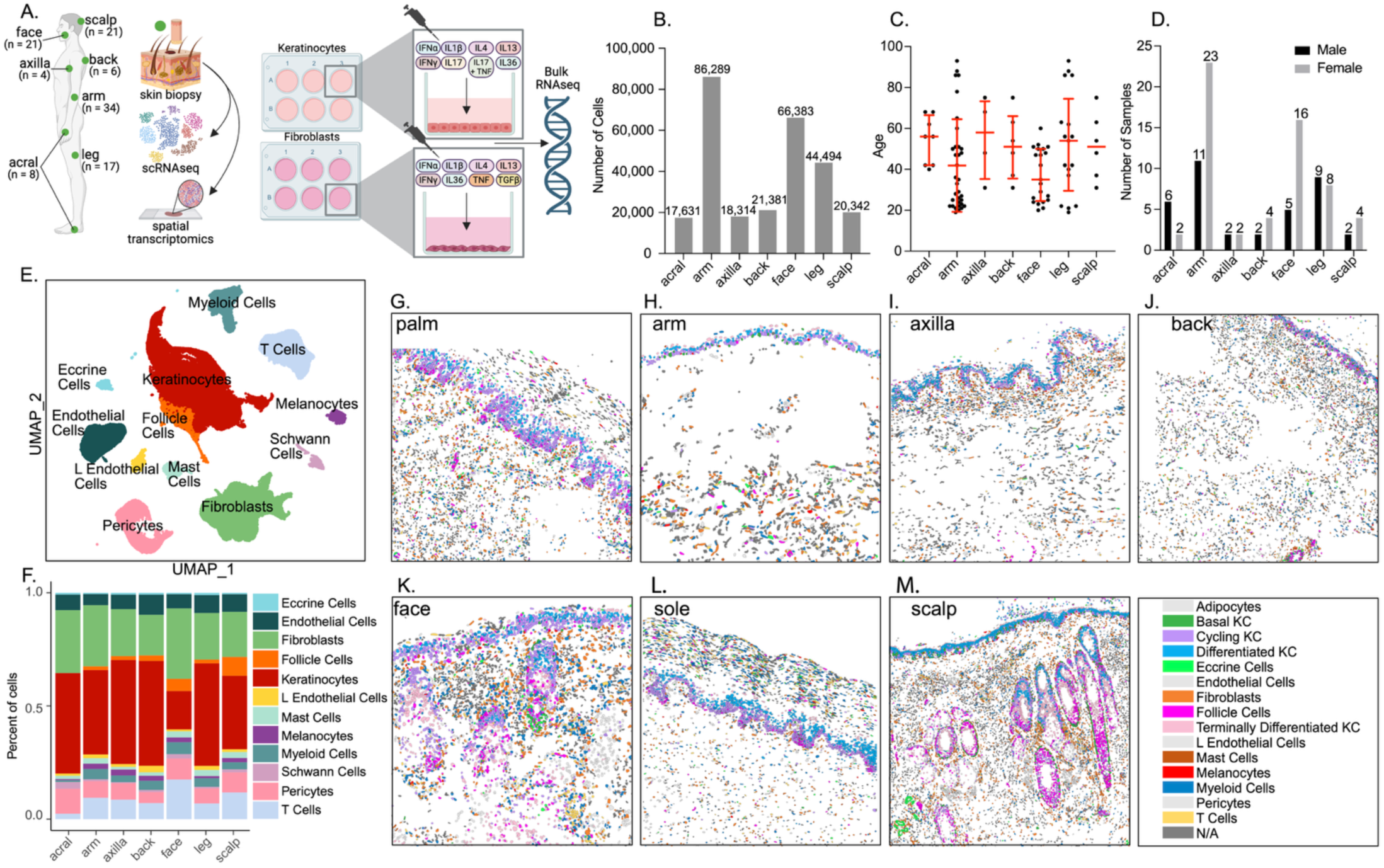
A single-cell atlas of healthy human skin reveals anatomical site-specific differences in cellular composition. (A) Schematic of experimental design and anatomical site sampling (B) Total number of high-quality cells captured per anatomical site following scRNA-seq and quality control (n = 17,831 cells from acral, n = 86,289 cells from arm, n = 18,314 cells from axilla, n = 21,381 cells from back, n = 66,383 cells from face, n = 44,494 cells from leg, and n = 20,342 cells from scalp) (C) Age distribution across anatomical sites (D) Sex distribution across anatomical sites (E) UMAP embedding of 274,834 cells colored by annotated cell types (F) Bar plot showing the relative abundance of major skin cell types across anatomical sites (G-M) CosMx spatial transcriptomics plots showing spatial distribution of cell types (Adipocytes, KC populations, Eccrine cells, Endothelial cells, Fibroblasts, Follicle cells, L Endothelial cells, Mast cells, Melanocytes, Myeloid cells, Pericytes and T cells) in palm, arm, axilla, back, face, sole and scalp skin.

### Anatomical site-specific KC subtypes exhibit distinct transcriptional and cytokine-response profiles in healthy human skin

To investigate the transcriptional diversity of KCs across anatomical sites, we performed subclustering analysis, which revealed distinct KC subpopulations with unique transcriptional signatures. Within the KC population, we identified basal KCs, cycling KCs, early differentiating KCs, differentiated KCs, terminally differentiated KCs, stress KCs, and hair follicle KCs (Fig. 2a). To define the molecular identity of these keratinocyte subpopulations, we examined key marker gene expression patterns. Basal KCs expressed high levels of *KRT14* and *KRT15*, and cycling KCs expressed high levels of *MKI67*, *TOP2A*, and *PCNA*, both consistent with their roles in early epidermal growth, proliferation and expansion (Fig. 2b). Early differentiating KCs expressed *DSG3* and *DSC3*, which have important roles in early epidermal differentiation (Fig. 2b). Differentiating KCs were enriched for *KRT1* and *KRT10*, and terminally differentiated KCs were enriched for *LOR* and *FLG2,* both indicative of terminal differentiation programs (Fig. 2b). Stress KCs uniquely displayed elevated expression of *KRT6A*, *KRT6C* and *KRT16*, suggesting increased involvement in stress-response and inflammatory signaling^25^ (Fig. 2b). Follicle cells highly expressed *KRT17,* which is important for hair follicle cycling, and *SOX9* and *LHX2*, which are important for maintenance of the stem cell niche within the follicle. The relative abundance of KC subsets also varied across anatomical sites. Face, scalp, and acral skin exhibited higher relative proportions of stress-responsive KCs, whereas arm, axilla, and leg skin contained greater proportions of early differentiating and fully differentiated KCs (Fig. 2c). Face and scalp skin also exhibited higher proportions of follicle cells relative to other anatomical sites. These differences highlight anatomical site-specific functional specialization, likely reflecting adaptations to environmental challenges, variations in hair follicle density, and distinct requirements for epidermal barrier integrity and immune regulation.

**Figure 2.**
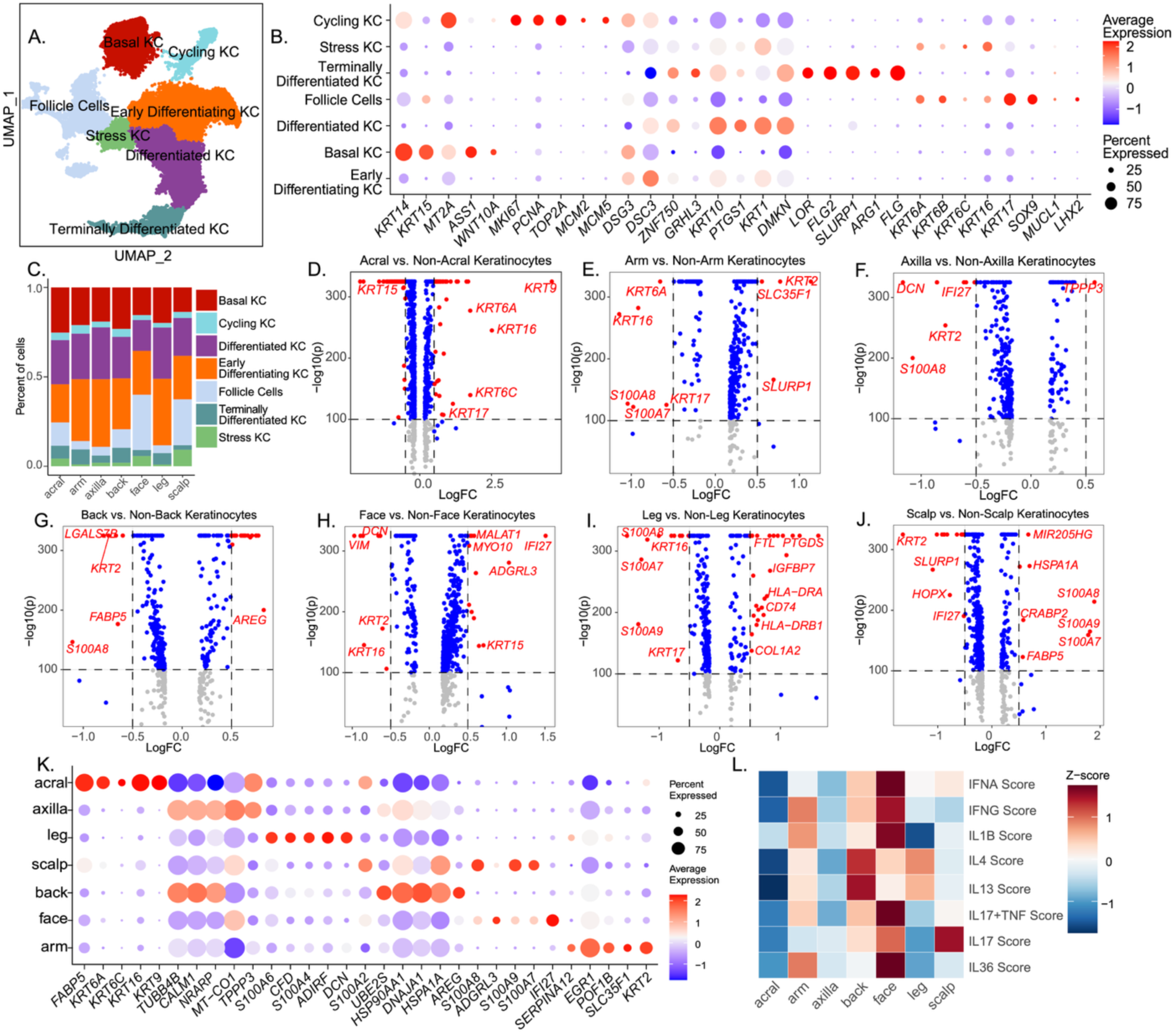
Anatomical site-specific KC subtypes exhibit distinct transcriptional and cytokine response profiles in healthy human skin. (A) UMAP visualization of KC subclusters across all anatomical sites (B) Dot plot showing marker gene expression used to define each KC subset (C) Bar plot showing the relative abundance of each KC subtype across anatomical sites (D–J) Volcano plots displaying differentially expressed genes in KCs from each anatomical site compared to all other sites (K) Dot plot showing the top 5 differentially expressed genes from KCs at each anatomical site (L) Heatmap of cytokine response module scores (IFN-α, IFN-y, IL-1β, IL-4, IL-13, IL-17, IL-17+TNF, IL-36) across KCs from different anatomical sites with data represented as a Z-score.

To explore site-specific transcriptional programs in KCs, we performed differential gene expression analysis across anatomical sites and generated volcano plots of significantly upregulated and downregulated genes in each anatomical site and a dotplot of the top 5 differentially expressed genes from each anatomical site (Fig. 2d-k, Supplementary Table S4). These data showed that Acral KCs displayed elevated expression of *KRT9*, as well as *KRT6A*, *KRT6C*, *KRT16*, and *KRT17*, which have been widely associated with the mechanical stress response^28^, whereas facial KCs demonstrated downregulation of *KRT16* and *KRT17*. In addition, facial KCs exhibited elevated expression of *IFI27*, a key interferon-stimulated gene (ISG), suggesting that increased expression of pro-inflammatory genes and decreased expression of stress-response genes may underlie the heightened susceptibility of facial skin to inflammatory skin diseases. Importantly, KCs from both facial and scalp skin exhibited high expression of the pro-inflammatory alarmin genes *S100A7*, *S100A8*, and *S100A9*, although to varying extents. While this may in part reflect the predisposition of these anatomical sites to inflammatory skin conditions such as scalp psoriasis or facial acne, it could also be influenced by the higher baseline UV exposure of the face, as these alarmins are known to be interferon-regulated genes induced by environmental triggers^29–31^. To assess signaling pathway activity, we performed Hallmark 50 pathway enrichment analysis across KCs from each anatomical site (Supplementary Fig. 2a). Facial and scalp KCs were highly active in terms of their signaling enrichment as compared to other anatomical sites. Facial KCs showed strong enrichment for “IL6-JAK-STAT3 signaling”, “inflammatory response”, and “TNF signaling via NFkB”, consistent with heightened immune signaling. Scalp KCs showed upregulation of hypoxia, oxidative phosphorylation, and xenobiotic metabolism, suggesting adaptation to environmental exposure and metabolic stress. To evaluate KC-to-KC interactions, we used CellChat to infer outgoing and incoming signaling strengths among KC subtypes (Supplementary Fig. 2b). Consistent with pathway enrichment, facial KCs also exhibited the highest incoming and outgoing interaction strengths relative to other anatomical sites.

Given the central role of KCs in inflammatory signaling, we next assessed cytokine response module scores across anatomical sites to evaluate regional variation in immune-related transcriptional programs. We generated module scores using differentially expressed gene lists generated bulk RNA-seq of N-TERT KCs stimulated with various cytokines including IFN-α, IFN-y, IL-1β, IL-4, IL-13, IL-17, IL-17+TNF, and IL-36 (Fig. 1a, Fig. 2l; Supplementary Table S5a-b). Strikingly, we observed the highest IFN-α, IFN-y, IL-1β, IL-17+TNF, and IL-36 module scores in facial KCs relative to other sites, indicating a highly inflammatory microenvironment, potentially driven by environmental insults such cumulative UV-exposure. In contrast, acral KCs demonstrated downregulation of all modules as compared to other anatomical sites, suggesting that their gene expression profiles may be more influenced by stress-response and metabolic cues resulting from mechanical forces, rather than immune-mediated signaling. In contrast, IL-4 and IL-13 module scores, which are associated with type 2 immune responses, are more prominent in back and leg KCs. Importantly, scalp KCs exhibited the highest IL17 score, suggesting that scalp has a bias towards Th17-associated signaling pathways and potential predisposition to diseases such as psoriasis. Hallmark 50 pathway analysis across anatomical sites also demonstrated that scalp is highly enriched for metabolic processes such as glycolysis and oxidative phosphorylation. Recent studies have demonstrated that the IL-17–HIF1α signaling axis contributes to epidermal metabolic dysfunction and plays a key role in sustaining inflammatory pathology in psoriasis^32^. Together, these findings show that KCs exhibit distinct transcriptional programs across different anatomical sites, with site-specific differences in differentiation, immune responses, and signaling networks. The differences in cytokine response scores suggest that regional variation in KC immune activation may contribute to site-specific predilection to inflammatory skin diseases such as psoriasis (IL-17-driven) and AD (IL-4/IL-13-driven).

### Anatomical site-specific fibroblast subtypes exhibit distinct transcriptional and cytokine response profiles in healthy human skin

To characterize the transcriptional diversity of fibroblasts across human skin, we performed subclustering analysis, identifying distinct fibroblast populations with site-specific transcriptional programs. The UMAP projection revealed fibroblast subsets including SFRP2^+^ fibroblasts, CCL19^+^ fibroblasts, TNN^+^ fibroblasts, CLDN1^+^ fibroblasts, FMO1^+^ fibroblasts, and FMO2^+^ fibroblasts (Fig. 3a). To confirm the identity and validate our annotation of these fibroblast subpopulations, we examined signature gene expression patterns using a dotplot (Fig. 3b). SFRP2^+^ fibroblasts expressed *SFRP2* and *MMP2* and *COL3A1*, key players of extracellular matrix (ECM) regulation, tissue homeostasis and regeneration^28,33,34^. CCL19^+^ fibroblasts, associated with immune modulation, exhibit high expression of *CCL19, CD74, TNFSF13B, APOE* and *C3*. TNN^+^ fibroblasts expressed *TNN*, *POSTN*, *COL11A1*, *CRABP1*, genes implicated in tissue remodeling and fibrosis^35–39^. CLDN1^+^ fibroblasts upregulated *CLDN1*, *SFRP4* and *NR2F2*, supporting their role in ECM remodeling^40,41^. FMO1^+^ and FMO2^+^ fibroblasts, enriched in metabolic activity, show elevated expression of *FMO1*, *FMO2*, *APOE*, and *FGFBP2*, suggesting roles in lipid and xenobiotic metabolism and stress responses^42,43^. Similar to KCs, we observed differences in abundance of fibroblast subsets across anatomical sites (Fig. 3c). Face and scalp fibroblasts contained the highest relative proportions of TNN^+^ fibroblasts compared to other anatomical sites. Similarly, they had relatively lower proportions of SFRP2^+^ fibroblasts. Conversely, acral fibroblasts skin had elevated proportions of FMO1^+^, FMO2^+^, and CLDN1^+^ fibroblasts. These findings suggest distinct functional specializations of fibroblast populations across anatomical sites, with facial and scalp skin exhibiting increased proportions of TNN^+^ fibroblasts, and decreased proportions of SFRP2^+^ fibroblasts. To spatially contextualize fibroblast subset distribution, we employed CosMX spatial transcriptomics, which revealed that these fibroblast populations occupy distinct dermal niches across anatomical sites (Supplementary Fig. 3a). For example, TNN⁺ fibroblasts were enriched in deeper dermal layers of facial and scalp skin, while FMO1⁺/FMO2⁺ fibroblasts were predominantly localized to acral skin. CCL19⁺ fibroblasts were often found near immune cell clusters in face and axilla, suggesting a more immune-interacting role.

**Figure 3.**
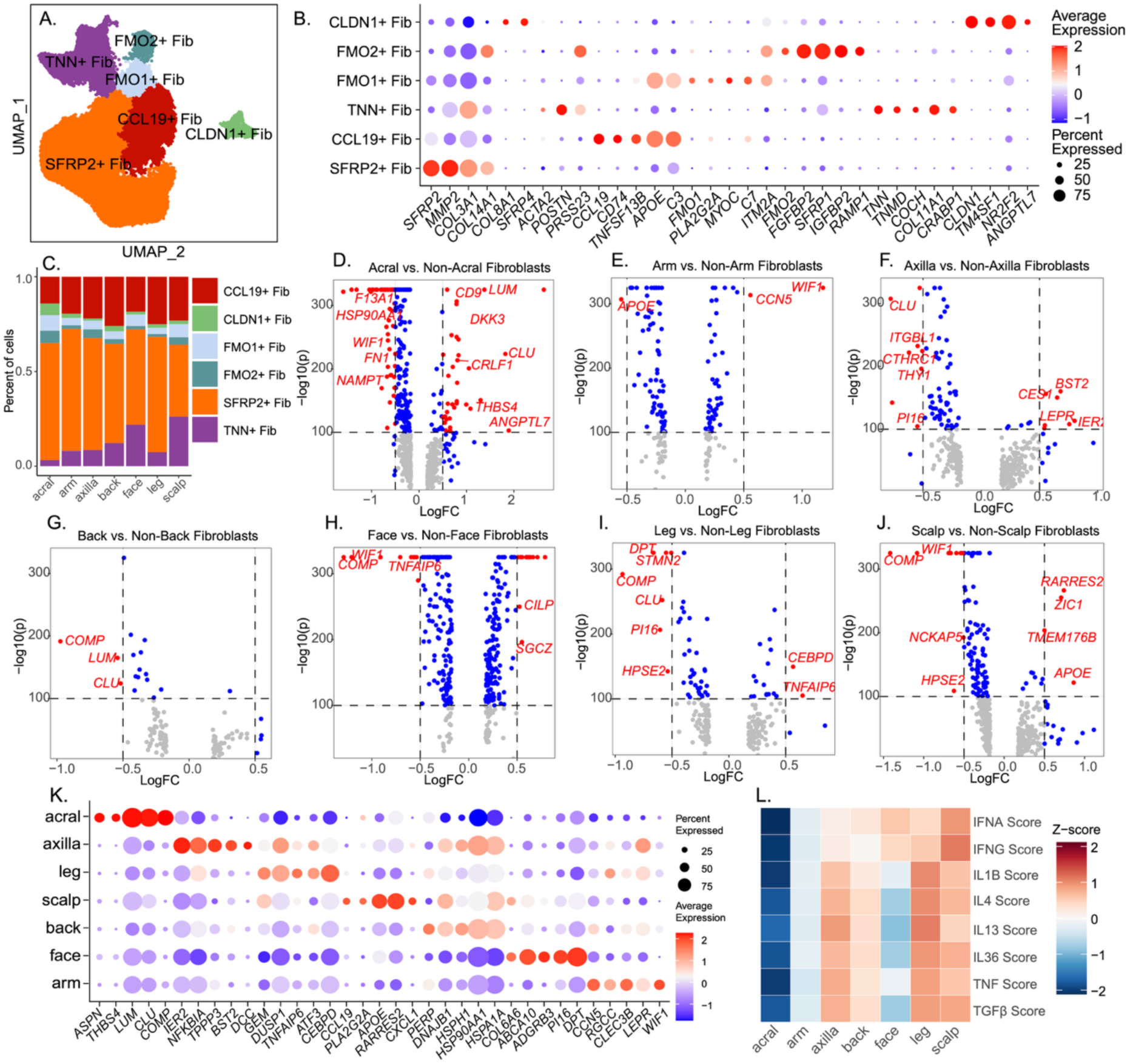
Anatomical site-specific fibroblast subtypes exhibit distinct transcriptional and cytokine response profiles in healthy human skin. (A) UMAP visualization of fibroblast subclusters across anatomical sites (B) Dot plot showing expression of representative marker genes for each fibroblast subtype (C) Bar plot showing the relative abundance of fibroblast subtypes across anatomical sites. (D–J) Volcano plots showing differentially expressed genes in fibroblasts from each anatomical site compared to all others. (K) Dot plot showing the top differentially expressed fibroblast markers across anatomical sites (L) Heatmap of cytokine response module scores (IFN-α, IFN-y, IL-1β, IL-4, IL-13, IL-36, TNF, TGFβ) across fibroblasts from different anatomical sites.

Differential gene expression analysis across fibroblasts identified significantly upregulated and downregulated genes in each body site (Fig. 3d-j, Supplementary Table S6). Acral fibroblasts exhibited upregulation of genes involved in ECM stability and ECM and tissue remodeling (*THBS4*, *ANGPTL7*), whereas facial fibroblasts showed elevated expression of genes related to tissue degeneration (*CILP*, *SGCZ*) and diminished expression of tissue development and repair genes (*COMP* and *TNFAIP6*). Axilla fibroblasts upregulated immune and metabolic markers (*CES1*, *BST2*, *IER2*). We next wanted to understand the functional roles of site-specific fibroblasts, so we performed Hallmark 50 pathway enrichment analysis (Supplementary Fig. 3b). We saw an interesting distribution of pathway enrichment with scalp fibroblasts demonstrating the highest propensity for inflammatory and fibrotic response, including “IFNG response”, “TNF signaling via NFkB”, “IL6/JAK/STAT3 signaling”, and “TGFβ signaling” relative to other anatomical sites. In contrast, facial fibroblasts were enriched for proliferative and UV-response pathways and acral fibroblasts were enriched for metabolic and stress-response pathways. Site-specific fibroblast signaling and communication analysis via CellChat revealed high fibroblast subset incoming and outgoing interactions in face and scalp fibroblasts as compared to other bodysites, consistent with what we observed in pathway enrichment (Supplementary Fig. 3c).

To investigate the immunomodulatory roles of fibroblasts across anatomical sites, we evaluated cytokine response module scores derived from differentially expressed gene lists generated through bulk RNA-seq of primary fibroblasts stimulated with various cytokines, including IFN-α, IFN-γ, IL-1β, IL-4, IL-13, IL-36, TNF, and TGFβ (Fig. 1a). This approach enabled us to assess site-specific patterns of immune signaling and modulation (Fig. 3l, Supplementary Table S7a-b). Module scores showed distinct inflammatory signatures in fibroblasts from different anatomical sites. Facial fibroblasts only exhibited elevated IFN-α and IFN-γ scores, suggesting enhanced antiviral responses or responses to chronic UV exposure, which is known to induce type I IFN signaling and S100 gene expression. In contrast, scalp fibroblasts exhibited high scores for all modules but especially IFN-y, IL-1β, IL-36, TNF and TGFβ scores, suggesting increased inflammatory activity and immune activation in scalp fibroblasts. This immune profile may contribute to the site-specific susceptibility of the scalp to fibroblast-driven diseases, including fibrosing alopecia and discoid lupus erythematosus (DLE). Leg and axilla fibroblasts demonstrated elevated scores for IL-1β, IL-4, IL-13, IL-36, TNF, and TGFβ modules. Similar to KCs, acral fibroblasts also had low scores across all modules. The differential immune signaling profiles suggest that fibroblasts actively shape tissue-specific inflammatory environments, potentially contributing to regional susceptibility to fibrosis and inflammatory skin disorders, for example as seen in discoid lupus erythematosus, a scarring subtype of lupus that is both associated with type I IFN signaling as well as fibrosis^44^. Collectively, these data emphasize specialized inflammatory adaptations of fibroblasts tailored to site-specific and environmental cues.

### Site-specific transcriptional programs in skin myeloid cells reveal regional immune states and enrichment of lupus-associated gene expression in facial skin

We next performed subclustering analysis on myeloid cells to explore their heterogeneity in healthy human skin and identified distinct myeloid subsets with site-specific transcriptional signatures. We identified Langerhans cells, conventional dendritic cell subsets (cDC1, cDC2a, cDC2b), plasmacytoid dendritic cells (pDCs), TREM2^+^ macrophages, M1 and M2 macrophages, and proliferating myeloid cells (Fig. 4a). To confirm the identity of these myeloid populations, we examined celltype-specific canonical gene expression signatures across subsets (Fig. 4b). Langerhans cells expressed high levels of *CD207*, and *CD1A*, consistent with their role in epidermal antigen presentation^45^ (Fig. 4b). cDC1 cells showed enrichment of *XCR1*, *WDFY4*, and *CLEC9A*, while cDC2a cells were characterized by expression of *CD1B* and *LAMP3*, and cDC2b cells by *CLEC10A* and *FCER1A*. These markers confirmed the distinct molecular profiles associated with their specialized roles in antigen presentation, with cDC1s efficiently presenting antigen to CD8 T cells and cDC2a and cDC2b’s presenting antigen to CD4 T cells^46,47^ (Fig. 4b). Macrophage subsets, including M1 and M2 macrophages, show differential expression of pro-inflammatory (*TNF*, *CD80*, and *CXCL10*) and anti-inflammatory (*MRC1*, *CD163*, and *SELENOP*) markers^48^ (Fig. 4b). These myeloid cell subsets also exhibited site-specific differences in their relative proportions, suggesting that skin site-specific immune surveillance and antigen presentation may be influenced by local environmental factors (Fig. 4c).

**Figure 4.**
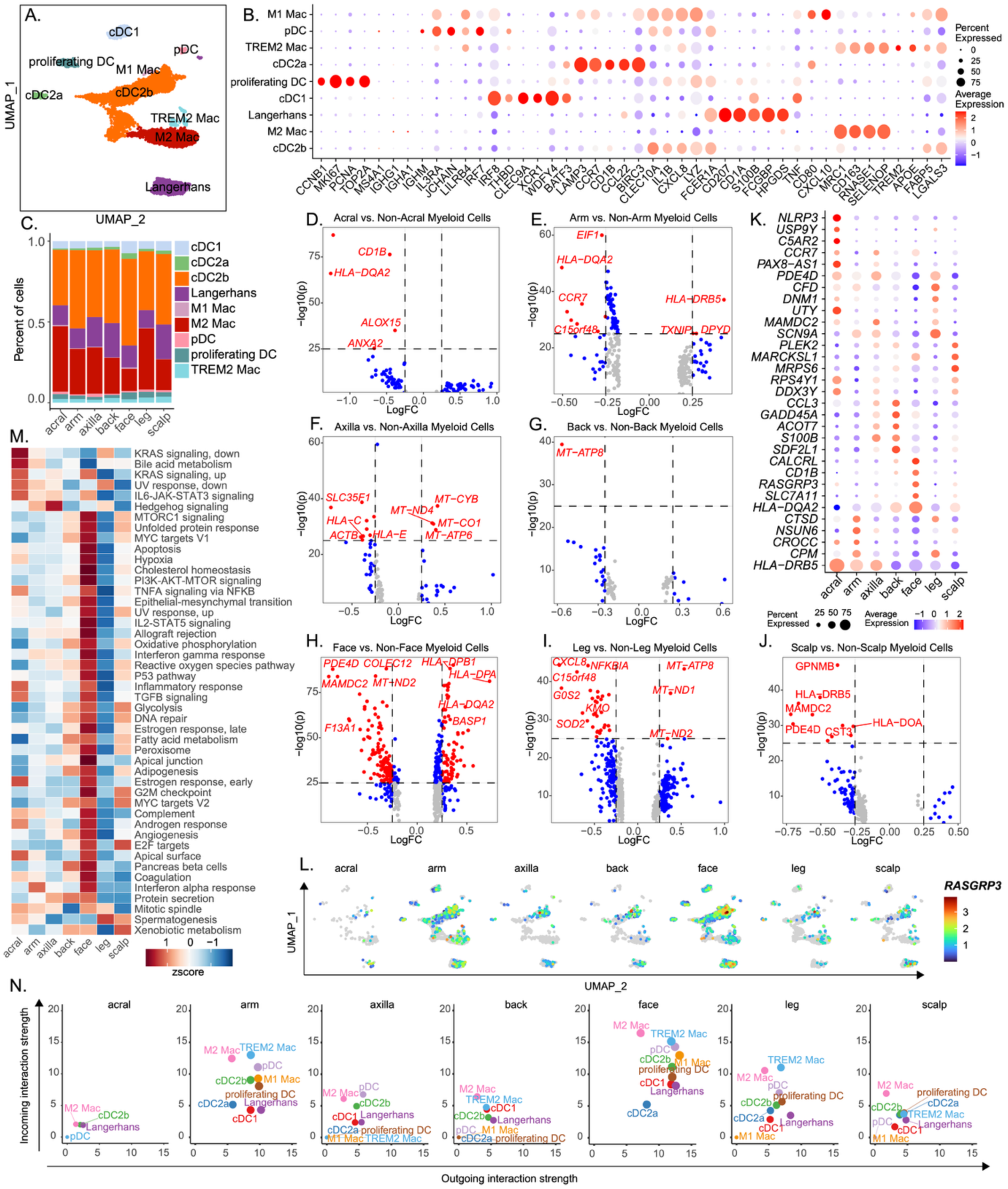
Site-specific transcriptional programs in skin myeloid cells reveal regional immune states and enrichment of lupus-associated gene expression in facial skin. (A) UMAP visualization of myeloid cell subclusters across anatomical sites (B) Dot plot showing expression of canonical marker genes across myeloid subtypes (C) Bar plot showing the relative abundance of each myeloid subset across anatomical sites (D–J) Volcano plots showing differentially expressed genes in myeloid cells from each anatomical site compared to all others. (K) Dot plot showing the top five most significant differentially expressed genes in myeloid cells from each anatomical site (L) Feature plot of *RASGRP3* expression across anatomical sites (M) Hallmark 50 enrichment heatmap showing pathway activity across myeloid cells from different anatomical sites (N) CellChat of predicted incoming and outgoing signaling interaction strength across myeloid subtypes and anatomical sites.

Differential gene expression analysis across anatomical sites identified significantly upregulated and downregulated genes specific to bodysite (Fig. 4d-j, Supplementary Table S8). Acral and back myeloid cells had fewer differentially expressed genes in comparison to other bodysites. Acral myeloid cells downregulated genes associated with antigen presentation and lipid metabolism (*ANXA2, HLA-DQA2, ALOX15, CD1B*) and back myeloid cells downregulated the mitochondrial metabolic gene, *MT-ATP8*. Arm myeloid cells exhibit increased expression of genes involved in immune cell migration, inflammatory responses, and metabolic adaptations (*TXNIP, DPYD*), along with decreased antigen presentation and protein translation-associated processes (*HLA-DQA2* and *EIF1*). Axilla myeloid cells uniquely upregulated mitochondrial genes (*MT-CYB, MT-CO1, MT-ND4, MT-ATP6*) and downregulated immunoregulatory genes (*HLA-E, HLA-C*), highlighting elevated metabolic activity. Facial myeloid cells were highly distinct from other sites based on their differential gene expression and showed increased expression of antigen-presentation genes (*HLA-DQA2, HLA-DPB1, HLA-DPA*) and downregulated extracellular matrix stabilization and remodeling genes (*F13A1, PDE4D*), suggesting active immune surveillance, coupled with downregulation of genes related to tissue repair. Leg and scalp cells notably downregulated inflammatory markers and certain antigen-presentation genes (*CXCL8*, *NFKBIA*, *HLA-DRB5*, and *HLA-DOA*), indicating reduced propensity for inflammatory responses. The top 5 differentially expressed genes identified the most significant site-specific transcriptional changes (Fig. 4k). Among these, RASGRP3, a gene previously implicated in SLE risk^49–51^, is significantly upregulated in facial myeloid cells. Feature plot analysis confirms its localized enrichment in facial myeloid cells, particularly in cDC subsets, suggesting a potential role in site-specific immune regulation (Fig. 4l). Given that RASGRP3 was linked to increased risk of facial rash in SLE^52^, these findings support the hypothesis that facial myeloid cells may play a unique role in lupus-associated skin manifestations.

To gain further insights into functional differences in myeloid cell states across anatomical sites, we performed Hallmark 50 pathway enrichment analysis on differentially expressed genes, revealing site-specific activation of immune and inflammatory pathways (Fig. 4m). Facial myeloid cells interestingly are most active in terms of pathway enrichment as compared to the other anatomical sites, showing most significant enrichment of several key pathways including inflammatory response pathways such as “interferon-gamma response”, “TNF signaling via NF-κB”. This suggested to us that facial myeloid cells are uniquely poised to signal and interact with other cell types to promote facial skin disease manifestations, which might explain facial predisposition of certain inflammatory skin diseases such as SLE and AD. Finally, we investigated myeloid-to-myeloid intercellular communication between myeloid subsets using CellChat and quantified incoming and outgoing interaction strengths within each anatomical site (Fig. 4n). Supporting of our observations with the hallmark pathway enrichment, facial myeloid populations, showed the highest interaction strengths overall, suggesting a regionally distinct inflammatory niche in facial skin. Together, these findings highlight the regional diversity of myeloid cell states, their distinct immune signaling programs across anatomical sites, and their unique inflammatory potential in facial skin, particularly through *RASGRP3* upregulation in facial myeloid subsets. Given the established link between *RASGRP3* genetic variants and SLE-related skin inflammation, these results suggest that site-specific immune regulation in facial myeloid cells may contribute to differential susceptibility to autoimmune skin manifestations.

### Regional specialization of T cell subsets in healthy human skin reveals distinct immune activation states and scalp-specific expression of psoriasis- and AD-associated genes

We were also interested in exploring the diversity of T cell populations in healthy human skin. T cell subsets included CD4 T cells, CD8 T cells, regulatory T cells (Tregs), tissue-resident memory T cells (Trms), naïve T cells, Natural killer T cells (NK-T cells) and gamma-delta T cells (GD-T cells) (Fig. 5a). To confirm the identity of these T cell subsets, we examined specific gene expression signatures across subsets (Fig. 5b). CD4 T cells expressed helper and homing markers (*CD4*, *IL7R*, *CCR7*), CD8 T cells exhibited cytotoxic signatures (*CD8A*, *CD8B*, *GZMA*, *GNLY*), and naive T cells expressed immature T cell genes (*CCR7*, *SELL*, *TCF7*). Trm cells expressed high levels of *CD69*, *CXCR6*, and *ITGAE*, indicative of long-term residency in the skin, and Tregs expressed *FOXP3*, *CTLA4* and *IL2RA*, defining their immune-modulatory function. NK-T cells exhibit strong expression of *KLRB1* and *NKG7*, consistent with their innate-like activation potential and gamma-delta T cells expressed innate-like receptors (*TRGV9*, *TRDC*). The different T cell subsets demonstrated site-specific variations in abundance, with scalp consisting of the highest percentage of CD4 T cells whereas face consisted of the highest percentage of Tregs (Fig. 5c). Interestingly, proportions of CD8 T cells were fairly uniform across anatomical sites, suggesting that T cell landscapes vary depending on anatomical site-specific functional requirements (Fig. 5c).

**Figure 5.**
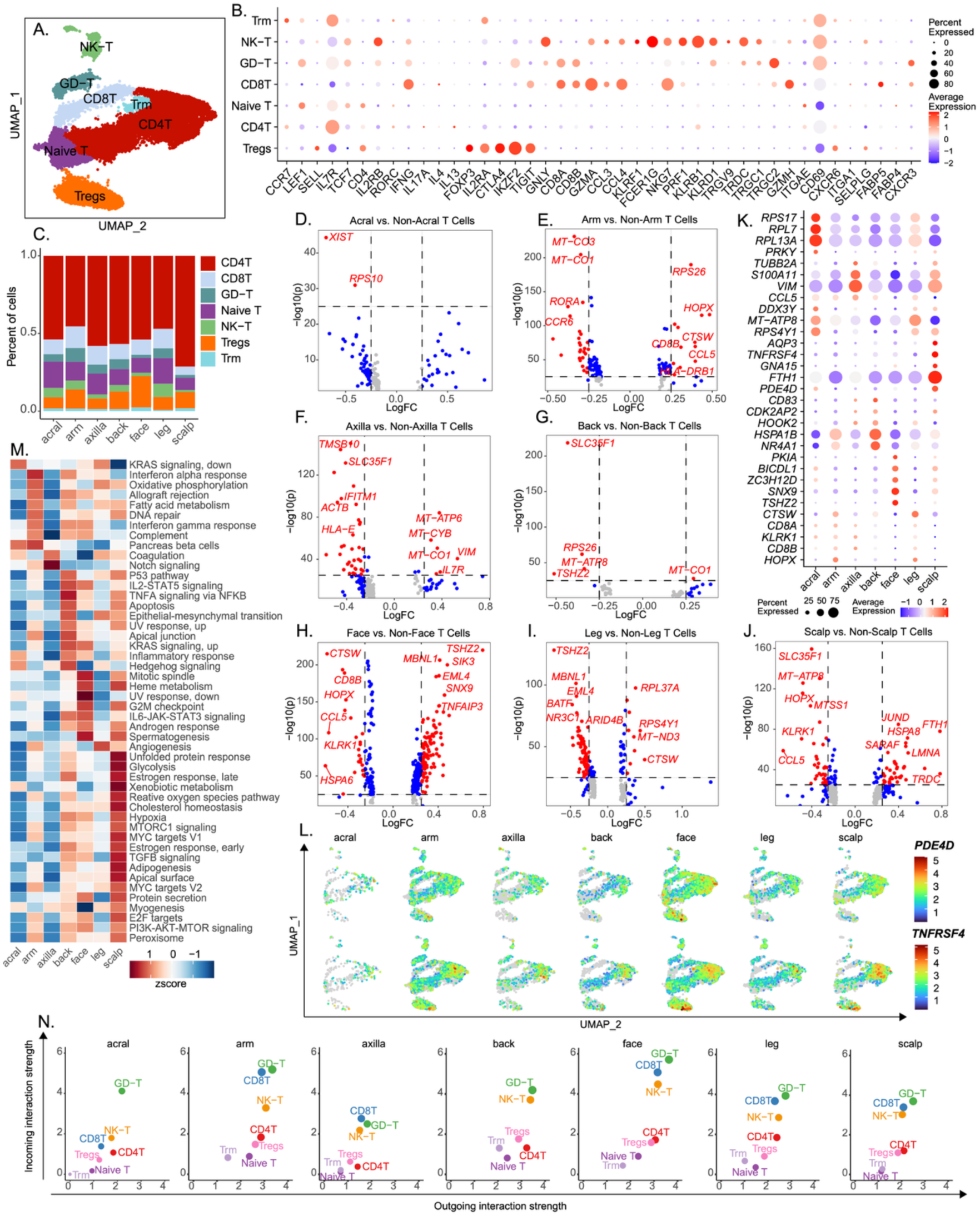
Regional specialization of T cell subsets in healthy human skin reveals distinct immune activation states and scalp-specific expression of psoriasis- and AD-associated genes. (A) UMAP visualization of T cell subclusters across anatomical sites (B) Dot plot showing expression of canonical marker genes defining each T cell subset (C) Bar plot showing the relative abundance of each T cell subset across anatomical sites (D–J) Volcano plots displaying differentially expressed genes in T cells from each anatomical site versus all others (K) Dot plot showing the top five most significant differentially expressed genes from each anatomical site (L) Feature plots showing selective enrichment of *PDE4D* and *TNFRSF4 (OX40)* in scalp CD4⁺ T cells and Tregs (M) Hallmark 50 pathway enrichment heatmap demonstrating site-specific pathway activation in T cells (N) CellChat analysis showing incoming and outgoing signaling interaction strengths across anatomical sites.

Differential gene expression across body sites (Fig. 5d-j, Supplementary Table S9) revealed that acral T cells were less distinct relative to T cells from other anatomical sites, as they exhibited relatively few differentially expressed genes. Arm T cells upregulated cytotoxic and metabolic genes (*HOPX, MT-CO1, CD8B*) while downregulating inflammatory markers (*HLA-DRB1, CCR6, CCL5*). Axilla and back T cells exhibited upregulation of mitochondrial and metabolic gene signatures (e.g., *MT-ATP6, MT-CYB, ACTB*) and axilla T cells also showed increased expression of T cell maintenance and homeostatic markers (*IL7R, VIM*). Facial T cells displayed increased expression of genes associated with cytotoxicity (*CTSW, CD8B*), inflammation and immune signaling (*CCL5, TNFAIP3*), and transcriptional regulation (*TSHZ2*), reflecting heightened immune activation. Leg T cells showed decreased expression of genes involved in transcriptional regulation and metabolism (*BATF, NR3C1*), while upregulating genes linked to translation (*RPL37A, RPS4Y1*) and cytotoxic functions (*CTSW*), suggesting a more regulatory or metabolic specialization compared to other sites. Scalp T cells had higher differential gene expression and exhibited elevated expression of genes associated with cellular stress, GD-T cell activity, and mitochondrial metabolism (*JUND, HSPA8, TRDC, FTH1, MT-ATP8*), along with decreased expression of cytotoxic, inflammatory, structural, and metabolic transport genes (*HOPX, KLRK1, CCL5, LMNA, SLC35F1*). This indicates specialized immune adaptations emphasizing stress response and metabolic activity, with reduced inflammatory and cytotoxic functions specific to scalp skin Notably, scalp T cells displayed distinct transcriptional patterns (Fig. 5k), with genes associated with *psoriasis* and AD showing site-specific upregulation. Scalp T cells showed significant upregulation of *PDE4D*, a gene implicated in psoriasis pathogenesis and targeted by PDE4 inhibitors for psoriasis treatment^53–55^. Additionally, *TNFRSF4* (OX40), a key player in type 2 immune responses and AD^56–59^, was also significantly upregulated in scalp T cells. Feature plot analysis confirmed localized enrichment of *PDE4D* and *TNFRSF4* in scalp CD4^+^ T cells and Tregs (Fig. 5l).

Hallmark 50 pathway enrichment analysis showed that scalp T cells demonstrated the highest enrichment of metabolic and stress-response pathways, such as “glycolysis”, “hypoxia”, “xenobiotic metabolism”, and “reactive oxygen species pathway”, suggesting that scalp T cells can adapt to a heightened state of metabolic activity and effectively respond to stress, enabling them to maintain immune surveillance (Fig. 5m). Interestingly, CellChat analysis revealed that face and arm T cells have the highest incoming and outgoing interaction strengths, and scalp T cells have comparatively lower interaction strengths, suggesting that scalp T cells rely on communication with other cell types to dynamically signal, regulate, and respond to inflammatory stimuli (Fig. 5n). Together, these findings demonstrate that T cells have different site-specific gene expression profiles and relative abundances of T cell subsets. Scalp T cells show significant upregulation of *PDE4D*, linked to psoriasis, and *TNFRSF4* encoding OX40 and linked to AD. The unique pathway activation profiles in different anatomical sites highlight the regional specialization of skin immunity, suggesting that therapeutic interventions targeting PDE4D or OX40 signaling may have differential effects across anatomical sites.

### Disease-specific upregulation of RASGRP3, PDE4D, and TNFRSF4 in SLE, psoriasis, and AD highlights their roles in anatomical site-specific skin inflammation

To investigate the role of *RASGRP3*, *PDE4D*, and *TNFRSF4* in inflammatory skin diseases, we analyzed an independent scRNA-seq dataset comparing control, SLE lesional and nonlesional skin, and psoriasis lesional and nonlesional skin biopsies. We identified the major skin cell types in this dataset and confirmed their identity and annotation using canonical cell type-specific gene expression signatures (Fig. 6a-b). We first probed for the mRNA expression of *RASGRP3, PDE4D, and TNFRSF4* in this dataset and observed that *RASGRP3* was significantly upregulated in both SLE lesional and psoriasis lesional skin as compared to control skin (Fig. 6c). Similarly, *PDE4D*, a phosphodiesterase linked to cyclic AMP signaling and inflammation, was also significantly increased in SLE and psoriasis lesional skin (Fig. 6d). *TNFRSF4* on the other hand was significantly upregulated in psoriasis lesional skin but not in SLE lesional skin (Fig. 6e), suggesting that TNFRSF4 has a stronger role in Th2/Th17-driven inflammation. Next, we performed immunohistochemistry (IHC) validation to confirm the expression of these markers at the protein level in healthy and diseased skin (Fig. 6f-h). IHC validation confirmed increased RASGRP3 protein levels in SLE lesional facial skin compared to healthy facial skin, further supporting it upregulation in lupus-associated skin inflammation. IHC also confirmed increased PDE4D protein levels in psoriatic scalp skin compared to healthy scalp skin, suggesting that PDE4D-driven inflammation may be particularly relevant in scalp lesions, potentially influencing treatment response to PDE4 inhibitors. Similarly, IHC showed increased TNFRSF4 protein levels in AD facial skin compared to healthy facial skin, confirming its association with Th2-driven immune activation. These results are consistent with previous studies identifying OX40 signaling as a therapeutic target in AD, while also suggesting a potential role in psoriasis pathogenesis. Together, these findings establish that RASGRP3, PDE4D, and TNFRSF4 exhibit disease-specific upregulation in SLE, psoriasis, and AD, reinforcing their potential roles in inflammatory skin diseases.

**Figure 6.**
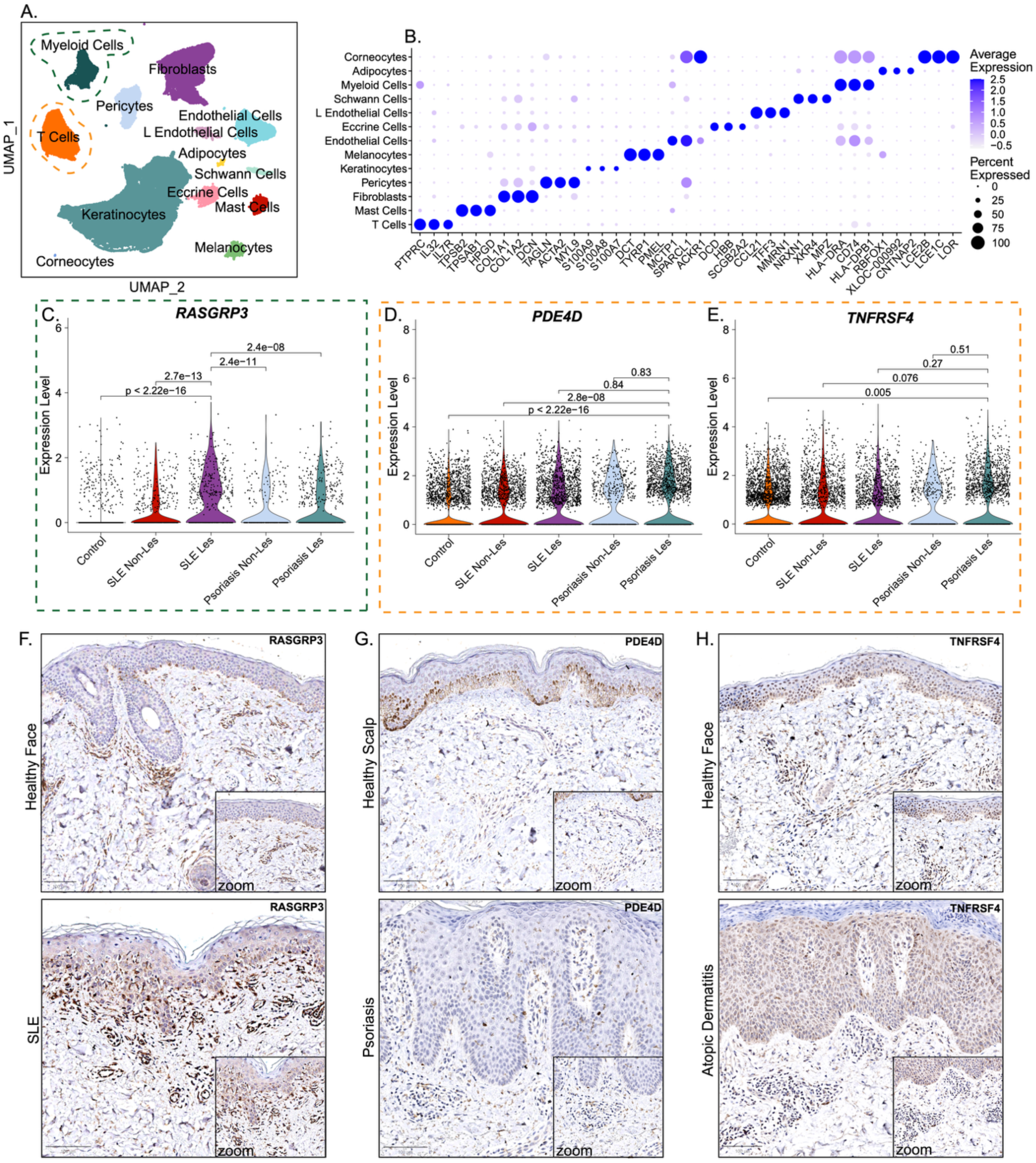
Disease-specific upregulation of RASGRP3, PDE4D, and TNFRSF4 in SLE, psoriasis, and AD highlights their roles in anatomical site-specific skin inflammation. (A) UMAP visualization of major skin cell types from an independent scRNA-seq dataset including control, SLE lesional and non-lesional skin, and psoriasis lesional and non-lesional skin (B) Dot plot showing expression of key marker genes across cell types in the inflammatory skin dataset (C–E) Violin plots of RASGRP3 (C), PDE4D (D), and TNFRSF4 (OX40) (E) expression across disease groups (F–H) Immunohistochemistry (IHC) validation (F) RASGRP3 protein expression in SLE lesional facial skin compared to healthy facial skin (G) PDE4D protein expression in psoriatic skin compared with healthy scalp skin (H) TNFRSF4 protein expression in AD facial skin compared with healthy facial skin.

### Genetic risk loci for inflammatory skin diseases align with site-specific gene expression programs, revealing potential mechanisms of regional disease susceptibility

To understand the relationship between genetic risk loci for inflammatory skin diseases and anatomical site-specific gene expression, we mapped genome-wide association study (GWAS) loci from acne, AD, SLE, and psoriasis to genes located within 200kb of significant GWAS signals. We analyzed the chromosomal distribution of mapped GWAS loci, which revealed distinct clustering patterns for each disease. Psoriasis-associated loci were dispersed across multiple chromosomes, while AD and acne loci were more concentrated in specific genomic regions (Fig. 7a-d). We then assessed the overlap of these disease-associated genes with differentially expressed genes across distinct anatomical sites in healthy skin, identifying regions where genetic risk loci coincide with transcriptional activity in specific anatomical sites. Our analysis revealed high variation in the number of GWAS-mapped genes overlapping with anatomical site-specific DEGs (Fig. 7e-f). Psoriasis- and acne-associated genes exhibited the most pronounced overlap with DEGs in scalp and facial skin, whereas SLE-associated loci were more evenly distributed across multiple anatomical sites, consistent with the systemic nature of lupus-related skin manifestations. Acne-associated loci were predominantly mapped to DEGs enriched in facial skin, reflecting the known predilection of acne lesions for sebaceous-rich areas. To explore anatomical site-specific expression of these mapped genes, we examined their expression levels across different anatomical sites, to identify distinct disease-specific and site-specific expression patterns (Fig. 7g). Several immune-related genes implicated in psoriasis and AD GWAS loci were most highly expressed in facial skin, reinforcing the role of regional immune variation in disease susceptibility. In contrast, SLE-associated genes were found across multiple anatomical sites, suggesting a broader systemic transcriptional signature rather than site-specific immune specialization. The expression of acne-associated genes in facial skin correlated with known sebaceous gland activity, further supporting the relevance of predisposing genetic factors in site-specific disease presentation. Together, these findings establish a direct link between genetic risk loci and regional gene expression programs in healthy skin, suggesting that pre-existing anatomical site variation in gene expression may contribute to site-specific disease susceptibility.

**Figure 7.**
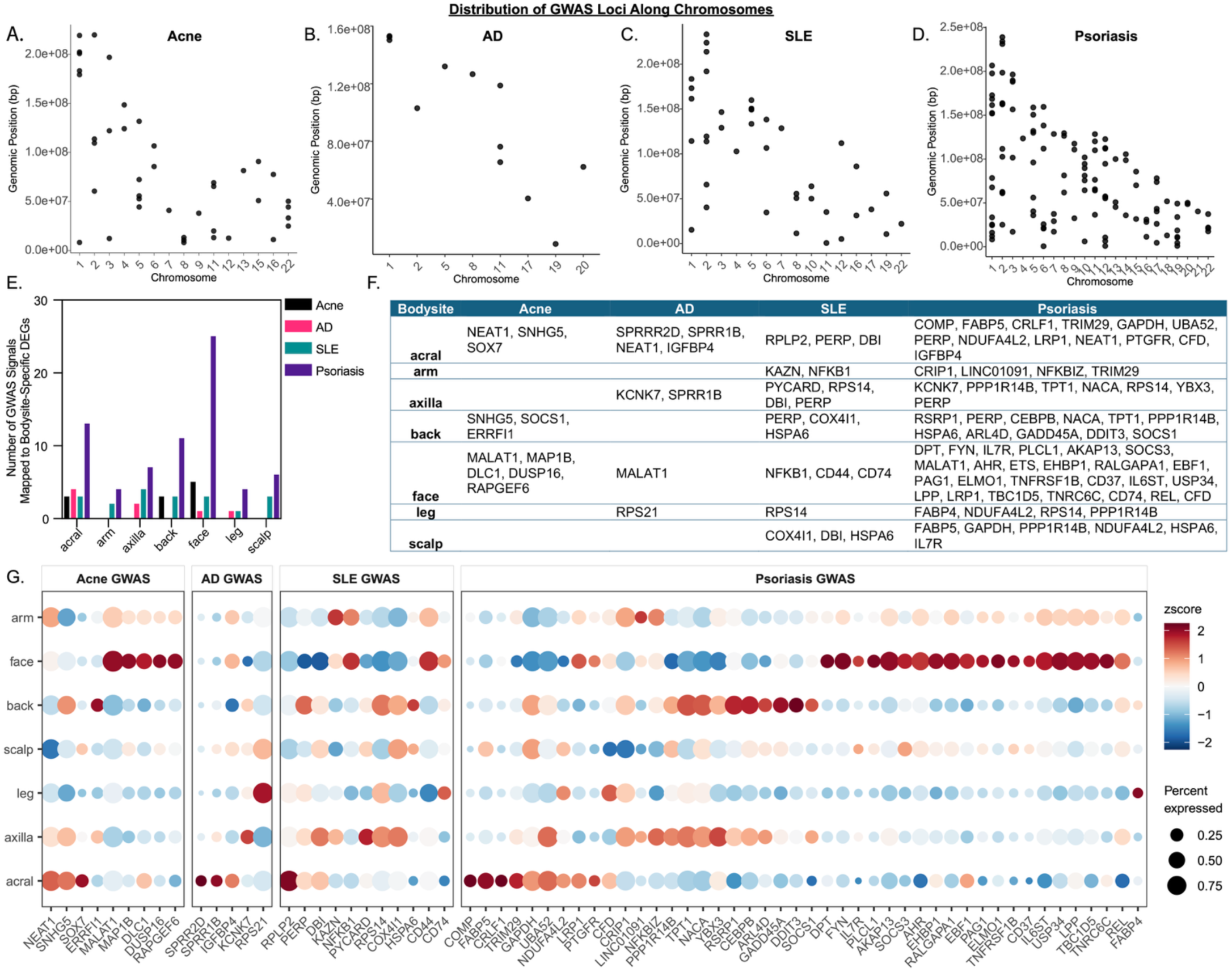
Genetic risk loci for inflammatory skin diseases align with site-specific gene expression programs, revealing potential mechanisms of regional disease susceptibility. (A–D) Chromosomal distribution of GWAS loci mapped to genes within ±200 kb of lead SNPs for (A) acne, (B) AD, (C) SLE and (D) psoriasis (E) Bar plot showing the number of GWAS-mapped genes overlapping with anatomical site-specific differentially expressed genes (F) Table displaying the disease-associated genes overlapping with anatomical site-specific DEGs across all anatomical sites (G) Dot plot showing the expression of selected GWAS-mapped genes across anatomical sites. Color indicates average scaled expression (z-score), and dot size represents the percentage of expressing cells.

## DISCUSSION

This study presents a high-resolution single-cell atlas of healthy human skin across different anatomical sites highlighting the transcriptional diversity and immune specialization of each site. Through detailed profiling of KCs, fibroblasts, myeloid cells, and T cells, we reveal how site-specific adaptations influence skin homeostasis and disease susceptibility. These findings provide a critical foundation for understanding regional immune regulation and its implications for explaining anatomical site predilection of inflammatory skin diseases such as SLE, psoriasis, and AD.

A key insight from our study is the regional and functional specialization of skin cell populations, particularly in immune surveillance and inflammatory signaling. KCs exhibited site-specific transcriptional programs, with facial KCs showing elevated expression of IL-17-related genes and enrichment of several pro-inflammatory pathways such as those involved in TNF signaling and interferon responses, suggesting an intrinsic inflammatory bias in sun-exposed skin. Similarly, scalp KCs had an elevated module score for IL-17-related genes but uniquely demonstrated metabolic pathways relating to glycolysis, oxidative phosphorylation and reactive oxygen species signaling. This regional priming may contribute to the site predilection observed in psoriasis and AD, which are driven by IL-17 and Th2-skewed immune responses, respectively, and also have a metabolic dysregulation component to disease pathogenesis^32,60–64^. Fibroblasts also demonstrated anatomical site-specific functional niches. Interestingly, scalp fibroblasts possessed a pronounced inflammatory profile, with elevated scores across all cytokine response modules. They also showed upregulation of pathways relating to inflammatory signaling and cellular metabolism. These findings suggest that scalp skin is highly metabolically active, and scalp fibroblasts might contribute to pro-inflammatory responses, while both scalp fibroblasts and KCs may have active roles in driving metabolic dysregulation. We also observed striking site-specific gene expression profiles among T cells and myeloid cells. Scalp T cells showed marked upregulation of *PDE4D*, a psoriasis susceptibility gene and target of PDE4 inhibitors (e.g., apremilast), implicating these cells as potential local drivers of inflammation. The OX40 receptor (*TNFRSF4*), known to be highly expressed in activated T cells from patients with AD and associated with Th2-driven inflammation, was also markedly elevated in scalp T cells, further supporting the scalp’s predisposition to heightened inflammatory immune activation^56,57^. In contrast, facial myeloid cells were enriched for *RASGRP3*, an gene linked with SLE risk and associated with malar rashes, a hallmark feature of disease, suggesting a potential mechanism for regional immune dysregulation in cutaneous manifestations of SLE^49,65^. Collectively, these findings underscore the importance of anatomical context in shaping skin immune gene regulation and disease susceptibility, with implications for targeted therapeutic strategies.

Our disease-focused analysis further validates the pathogenic relevance of *RASGRP3*, *PDE4D*, and *TNFRSF4* in inflammatory skin disorders. *RASGRP3* is upregulated in SLE lesional skin with genetic variants in *RASGRP3* linked to SLE susceptibility,^50–52^ and its enrichment in facial myeloid cells suggests a potential molecular link between genetic risk and localized skin inflammation in SLE. Similarly, our analysis revealed that *PDE4D* expression is highly elevated in psoriatic lesional skin, particularly within T cells, aligning with its known enrichment in the dermal inflammatory infiltrate of psoriasis lesions and further underscore its clinical relevance as a therapeutic target of PDE4 inhibitors^66^. Likewise, *TNFRSF4* (OX40), a key regulator of T cell activation, is significantly enriched in psoriatic lesions, suggesting that OX40 blockade, a current treatment modality for AD, may also hold potential for psoriasis treatment. These findings highlight the importance of regional immune specialization in the skin, highlight potential biomarkers of site-specific disease susceptibility, and reinforce the concept that different anatomical sites may have distinct inflammatory thresholds and therapeutic responses. Avenues for further investigation include determining how environmental factors such as UV exposure, microbiome composition, and barrier integrity contribute to site-specific immune programming, potentially influencing disease predisposition, progression, and treatment response.

While this study provides a large and high-powered spatial and single-cell datasets to explore numerous questions on skin homeostasis and site-specific predilections for skin disease, some limitations should be acknowledged. Our analysis represents a static transcriptional snapshot, and further longitudinal studies are needed to determine how these site-specific immune programs evolve over time in response to environmental triggers and disease progression. Additionally, future investigations incorporating functional assays, *ex vivo* skin models, and *in vivo* validation studies will be essential to delineate causal mechanisms linking regional immune dysregulation to disease pathogenesis. In conclusion, this study provides a large single-cell atlas of healthy human skin, revealing site-specific adaptations in KCs, fibroblasts, myeloid cells, and T cells that influence immune homeostasis and disease susceptibility. Our findings demonstrate that regional immune landscapes shape both physiological and pathological processes, with direct implications for SLE, psoriasis, and AD. They also open the possibility for precision dermatology approaches, emphasizing the importance of targeting site-specific immune pathways for more effective therapeutic interventions. Importantly, this atlas serves as a foundational resource for the research community, enabling future studies to build upon these insights and further dissect the regional complexity of human skin immunity.

## METHODS

### Human samples and IRB statement

The study was approved by the University of Michigan Institutional Review Board (IRB) (HUM00174864), and all patients provided written informed consent. Eligibility criteria included healthy and normal appearing skin in patients without any inflammatory skin disease diagnosis.

### scRNA-seq library preparation, sequencing, and alignment

This scRNA-seq dataset was generated with 96 healthy skin biopsies from seven different anatomical sites, including acral, arm, axilla, back, face, leg, and scalp. Single-cell suspensions for scRNA-seq were generated by first incubating skin biopsies overnight at 4°C in 0.4% dispase (Life Technologies) prepared in Hank’s Balanced Saline Solution (Gibco). The epidermis and dermis were then separated. Epidermal tissue was digested with 0.25% Trypsin-EDTA (Gibco) and 10 U/mL DNase I (Thermo Scientific) for 1 hour at 37°C, followed by quenching with FBS (Atlanta Biologicals) and filtration through a 70μm mesh. Dermal tissue was minced and digested in a solution containing 0.2% Collagenase II (Life Technologies) and 0.2% Collagenase V (Sigma) in plain medium for 1.5 hours at 37°C, then filtered through a 70μm mesh. Epidermal and dermal cell suspensions were combined in a 1:1 ratio. Library preparation was performed at the University of Michigan Advanced Genomics Core using the 10X Chromium system with v2 and v3 chemistry. The libraries were then sequenced on the Illumina NovaSeq 6000 platform, generating 150 bp paired-end reads. Data processing, including quality control, read alignment to hg38, and gene quantification, was performed using the 10X Cell Ranger software. Finally, the samples were merged into a single expression matrix using the cellranger aggr pipeline.

### Cell clustering and cell type annotation

A total of 274,834 cells were sequenced and data was processed with Seurat v3.0. Quality control measures were applied to remove low-quality cells, excluding cells with fewer than 200 genes, more than 7,500 genes, or mitochondrial gene transcripts exceeding 20%. Ambient RNA was removed using the R package SoupX v1.6.2 and doublets were removed using R package scDblFinder v1.12.0. The data was normalized using the NormalizeData function and the “LogNormalize” method. The data was scaled using the ScaleData function and the top 2000 most variable genes were identified using the FindVariableFeatures function. Dimensionality reduction was performed using RunPCA on the 2000 most variable genes to derive principal components (PCs) and Uniform Manifold Approximation and Projection (UMAP) was applied to the top 30 PCs. Batch effect correction was conducted using R package Harmony v1.0, with donor as the batch variable. Following batch correction, cells were clustered using the FindNeighbors function with shared nearest neighbor (SNN) modularity optimization-based clustering. Then the FindClusters function was performed for modularity optimization-based clustering with a resolution of 0.5 to obtain biologically meaningful clusters. For visualization, the RunUMAP function was applied to the top 30 principal components to generate a Uniform Manifold Approximation and Projection (UMAP) representation of the data. Cluster-specific marker genes were identified using the FindAllMarkers function in Seurat v4.4.1. Cell type assignments were determined by comparing these marker genes to curated cell-type signature genes.

### Cell type subclustering

Sub-clustering was conducted on the predominant cell types using the same methods described above. In brief, data normalization was conducted using Seurat’s NormalizeData function, highly variable features were identified with the FindVariableFeatures function (selection method: “vst”), and all genes were used for scaling with ScaleData. Principal Component Analysis (PCA) was performed with 50 components using RunPCA. To correct for batch effects, Harmony integration was applied using RunHarmony, with donor as the batch variable and a maximum of 50 iterations. The integrated dataset was used for neighborhood graph construction (FindNeighbors) and clustering (FindClusters), with an initial resolution of 1.0. UMAP was performed using RunUMAP based on Harmony-reduced dimensions. A second round of neighbor detection and clustering was performed using UMAP-reduced dimensions with a lower resolution of 0.1 to refine subcluster identification. Subclusters characterized primarily by mitochondrial gene expression, indicative of low-quality cells, were excluded from further analysis. Cell subtypes were assigned by comparing the marker genes of each subcluster with established canonical subtype signature genes.

### Downstream scRNA-seq analyses

All downstream analyses were performed in R (v4.4.1) using the Seurat package (v4.3.0) and custom visualization tools from the scCustomize^67^, SeuratExtend^68^, and dittoSeq^69^ packages. We visualized cell type distributions using DimPlot_scCustom for UMAP-based clustering and dittoBarPlot to quantify the relative abundance of each myeloid subset across anatomical anatomical sites. Colors were manually curated for consistent visualization across plots. To identify marker genes, we used Seurat’s FindAllMarkers and FindMarkers functions with the Wilcoxon rank-sum test (test.use = “wilcox”), using default multiple testing correction and a minimum log fold-change threshold of 0.5. We applied filters to remove low-confidence or uninformative genes, including predicted genes (e.g., AL, AC, AP), keratin genes (KRT), and long intergenic noncoding RNAs (LINC). Top markers were extracted using top_n by average log2FoldChange within each group. Volcano plots were generated for each anatomical site versus all others using a custom VolcanoPlot() function, highlighting the top 50 differentially expressed genes (adjusted *p* < 0.05, logFC > 0.25). We generated dot plots of canonical cell type markers using DotPlot_scCustom() and DotPlot2(), with expression scaled and colored from blue (low) to red (high). We calculated cytokine-specific module scores using gene sets derived from in vitro stimulation experiments previously published by Billi et al., 2022^24^. Module scores were computed using the AddModuleScore function for each cytokine gene set. Scores were appended to the seurat object, renamed (e.g., IFNΑ_Score), and visualized using violin plots grouped by anatomical site. Mean score values were shown for each group, and statistical comparisons were performed using the Wilcoxon rank-sum test. To assess pathway-level differences, we performed single-cell gene set analysis using AUCell-based scoring via the GeneSetAnalysis() function with the MSigDB Hallmark 50 gene set. Pathway enrichment statistics were computed using CalcStats grouped by anatomical site and visualized with a custom Heatmap function showing z-scored activity. We performed gene-specific expression comparisons (RASGRP3, PDE4D, TNFRSF4) expression across both cell types and anatomical sites using split violin plots (VlnPlot2), applying Wilcoxon rank-sum tests to compare expression between anatomical sites, with significance denoted using adjusted *p*-values. To compare intercellular communication roles across skin anatomical sites, we performed signaling role analysis using the CellChat package in R. In brief, we processed our Seurat object according to the CellChat vignette^70^ and generated CellChat objects for each anatomical site (acral, arm, axilla, back, face, leg, and scalp). We then compiled the individual CellChat objects into a list and merged using the mergeCellChat and updated the merged object using updateCellChat to harmonize metadata and signaling networks across datasets. To evaluate the communication role of each signaling pathway, we generated scatterplots using netAnalysis_signalingRole_scatter, which display incoming and outgoing interaction strengths. Separate plots were generated for each anatomical site to reveal region-specific signaling dynamics.

### Mapping GWAS Signals to anatomical site-specific differentially expressed genes

Genome-wide association study (GWAS) data for acne-, AD-, SLE-, and Psoriasis-associated loci were imported and processed in R. Genomic positions of significant SNPs were extracted and mapped to nearby genes using the biomaRt package by querying the Ensembl database (GRCh38). For each SNP, protein-coding genes located within a ±200 kb window were identified. Unique gene symbols were assigned to each locus and appended to the dataset. To enable visualization of locus distribution, a scatter plot of SNP positions across chromosomes was generated using ggplot2. Genes mapped to multiple loci were separated into individual rows using tidyr, and empty or ambiguous entries were filtered out to generate a cleaned dataset. Genes were also grouped by chromosome and summarized into a reference table.

### Spatial transcriptomics

Flat files (exprMat_file.csv.gz, fov_positions_file.csv.gz, metadata_file.csv.gz, polygons.csv.gz, tx_file.csv.gz) were exported from AtoMX v1.3.2, the Spatial Informatics platform of Nanostring. During preprocessing, cells with fewer than 20 counts, negative probes (feature names starting with “Neg”), and system probes (feature names starting with “System”) were removed from analysis. Cell type annotations were performed using the R package InsituType v1.0.0 in a supervised way, utilizing a reference profile from single-cell RNA sequencing dataset. With the R package Giotto v4.1.0, a Giotto object was created based on the flat files. Information including the gene expression matrix, cell polygons, and spatial locations of transcripts for each field of view (FOV), were incorporated into a sub-Giotto object with the Giotto function “createGiottoObjectSubcellular”. The sub-Giotto objects of all FOVs were then merged to a comprehensive Giotto object using the Giotto function “joinGiottoObjects”. Insitu plots showing the distribution of selected features overlapped with cells were generated using the Giotto function “spatInSituPlotPoints”.

### Immunohistochemistry

Paraffin-embedded skin sections from excisional biopsies of healthy control, SLE, psoriasis, and AD patients were baked at 60 °C for 30 minutes, deparaffinized, and rehydrated through graded alcohols^22^. Antigen retrieval was performed in pH 9 buffer using a pressure cooker at 125 °C for 30 seconds. Following cooling, sections were treated with 3% hydrogen peroxide for 5 minutes to quench endogenous peroxidase activity, then blocked with 10% goat serum for 30 minutes at room temperature^22^. Primary antibodies were applied overnight at 4 °C, including: RASGRP3 (ThermoFisher Scientific, MA5-26504; 1:100), PDE4D (Millipore Sigma, HPA045895; 1:100), TNFRSF4/CD134 (ThermoFisher Scientific, BS-2685R; 1:100), and Goat IgG isotype control (Jackson ImmunoResearch, 005-000-003)^22^. The next day, slides were washed, incubated with appropriate HRP-conjugated goatαrabbit secondary antibody for 30 minutes, and developed using diaminobenzidine (DAB) substrate prior to imaging^22^.

## ACKNOWLEDGEMENTS

This work was supported by the National Institute of Health: P30-AR075043 (J.E.G.).

## AUTHOR INFORMATION

Johann E. Gudjonsson supervised this work.

### Contributions

Conceptualization – S.M., A.C.B., J.E.G.

Study materials and resources – P.H., J.H.K., K.G., J.M.K., M.N., R.L.M., L.C.T.

Methods and investigation – S.M., R.B., Y.G., X.X., J.F., J.K. Writing – S.M., J.E.G

Critical feedback – A.C.B., B.K., V.V.D., J.M.K., R.B., M.K.S., M.N., K.G., V.W.R., S.N., B.A.

## REFERENCES

1 Damiani, G. et al. The Global, Regional, and National Burden of Psoriasis: Results and Insights From the Global Burden of Disease 2019 Study. Frontiers in Medicine 8 (2021). 10.3389/fmed.2021.743180

2 Izmirly, P. M. et al. Prevalence of Systemic Lupus Erythematosus in the United States: Estimates From a Meta-Analysis of the Centers for Disease Control and Prevention National Lupus Registries. Arthritis & Rheumatology 73, 991–996 (2021). 10.1002/art.41632

3 Lee, H. J., Hong, Y. J., Han, K. D. & Lee, J. H. Atopic Dermatitis Severity and Risk for Psoriasis: A Nationwide Population-Based Study. Dermatology 240, 262–270 (2024). 10.1159/000536143

4 Tian, J. et al. Global epidemiology of atopic dermatitis: a comprehensive systematic analysis and modelling study. Br J Dermatol 190, 55–61 (2023). 10.1093/bjd/ljad339

5 Ali, Z. et al. Assessing anatomical distribution of atopic dermatitis identifies a cluster of patients with late onset and low risk of asthma and allergy: An observational study. Health Sci Rep 6, e1219 (2023). 10.1002/hsr2.1219

6 Dopytalska, K., Sobolewski, P., Blaszczak, A., Szymanska, E. & Walecka, I. Psoriasis in special localizations. Reumatologia 56, 392–398 (2018). 10.5114/reum.2018.80718

7 Kole, A. K. & Ghosh, A. Cutaneous manifestations of systemic lupus erythematosus in a tertiary referral center. Indian J Dermatol 54, 132–136 (2009). 10.4103/0019-5154.53189

8 Stull, C., Sprow, G. & Werth, V. P. Cutaneous Involvement in Systemic Lupus Erythematosus: A Review for the Rheumatologist. J Rheumatol 50, 27–35 (2023). 10.3899/jrheum.220089

9 Vale, E. & Garcia, L. C. Cutaneous lupus erythematosus: a review of etiopathogenic, clinical, diagnostic and therapeutic aspects. An Bras Dermatol 98, 355–372 (2023). 10.1016/j.abd.2022.09.005

10 Weidinger, S., Beck, L. A., Bieber, T., Kabashima, K. & Irvine, A. D. Atopic dermatitis. Nat Rev Dis Primers 4, 1 (2018). 10.1038/s41572-018-0001-z

11 Yan, B. X. et al. Cutaneous and Systemic Psoriasis: Classifications and Classification for the Distinction. Front Med (Lausanne) 8, 649408 (2021). 10.3389/fmed.2021.649408

12 Jiao, Q. et al. Skin homeostasis: Mechanism and influencing factors. Journal of Cosmetic Dermatology 23, 1518–1526 (2024). 10.1111/jocd.16155

13 Yousef, H., Alhajj, M., Fakoya, A. O. & Sharma, S. in StatPearls (2025).

14 Nemes, Z. & Steinert, P. M. Bricks and mortar of the epidermal barrier. Experimental & Molecular Medicine 31, 5–19 (1999). 10.1038/emm.1999.2

15 Uitto, J., Olsen, D. R. & Fazio, M. J. Extracellular matrix of the skin: 50 years of progress. J Invest Dermatol 92, 61S–77S (1989). 10.1111/1523-1747.ep13075039

16 Belkaid, Y. & Segre, J. A. Dialogue between skin microbiota and immunity. Science 346, 954–959 (2014). 10.1126/science.1260144

17 Byrd, A. L., Belkaid, Y. & Segre, J. A. The human skin microbiome. Nature Reviews Microbiology 16, 143–155 (2018). 10.1038/nrmicro.2017.157

18 Reynolds, G. et al. Developmental cell programs are co-opted in inflammatory skin disease. Science 371, eaba6500 (2021). 10.1126/science.aba6500

19 Costello, C. M., Pittelkow, M. R. & Mangold, A. R. Acral Melanoma and Mechanical Stress on the Plantar Surface of the Foot. New England Journal of Medicine 377, 395–396 (2017). 10.1056/nejmc1706162

20 D’Orazio, J., Jarrett, S., Amaro-Ortiz, A. & Scott, T. UV Radiation and the Skin. International Journal of Molecular Sciences 14, 12222–12248 (2013). 10.3390/ijms140612222

21 He, H. et al. Single-cell transcriptome analysis of human skin identifies novel fibroblast subpopulation and enrichment of immune subsets in atopic dermatitis. J Allergy Clin Immunol 145, 1615–1628 (2020). 10.1016/j.jaci.2020.01.042

22 Ma, F. et al. Single cell and spatial sequencing define processes by which keratinocytes and fibroblasts amplify inflammatory responses in psoriasis. Nature Communications 14 (2023). 10.1038/s41467-023-39020-4

23 Perez, R. K. et al. Single-cell RNA-seq reveals cell type-specific molecular and genetic associations to lupus. Science 376, eabf1970 (2022). 10.1126/science.abf1970

24 Billi, A. C. et al. Nonlesional lupus skin contributes to inflammatory education of myeloid cells and primes for cutaneous inflammation. Science Translational Medicine 14 (2022). 10.1126/scitranslmed.abn2263

25 Cohen, E. et al. Significance of stress keratin expression in normal and diseased epithelia. iScience 27, 108805 (2024). 10.1016/j.isci.2024.108805

26 Ma, F. et al. Systems-based identification of the Hippo pathway for promoting fibrotic mesenchymal differentiation in systemic sclerosis. Nature Communications 15 (2024). 10.1038/s41467-023-44645-6

27 Van Straalen, K. R. et al. Single-cell sequencing reveals Hippo signaling as a driver of fibrosis in hidradenitis suppurativa. Journal of Clinical Investigation 134 (2024). 10.1172/jci169225

28 Bonnans, C., Chou, J. & Werb, Z. Remodelling the extracellular matrix in development and disease. Nature Reviews Molecular Cell Biology 15, 786–801 (2014). 10.1038/nrm3904

29 Grimbaldeston, M. A., Geczy, C. L., Tedla, N., Finlay-Jones, J. J. & Hart, P. H. S100A8 Induction in Keratinocytes by Ultraviolet A Irradiation Is Dependent on Reactive Oxygen Intermediates. Journal of Investigative Dermatology 121, 1168–1174 (2003). 10.1046/j.1523-1747.2003.12561.x

30 Harder, J., Tsuruta, D., Murakami, M. & Kurokawa, I. What is the role of antimicrobial peptides (AMP) in acne vulgaris? Exp Dermatol 22, 386–391 (2013). 10.1111/exd.12159

31 Wang, S. et al. S100A8/A9 in Inflammation. Frontiers in Immunology 9 (2018). 10.3389/fimmu.2018.01298

32 Subudhi, I. et al. Metabolic coordination between skin epithelium and type 17 immunity sustains chronic skin inflammation. Immunity 57, 1665–1680 e1667 (2024). 10.1016/j.immuni.2024.04.022

33 Lin, H. et al. sFRP2 activates Wnt/β-catenin signaling in cardiac fibroblasts: differential roles in cell growth, energy metabolism, and extracellular matrix remodeling. American Journal of Physiology-Cell Physiology 311, C710–C719 (2016). 10.1152/ajpcell.00137.2016

34 Tabib, T. et al. Myofibroblast transcriptome indicates SFRP2hi fibroblast progenitors in systemic sclerosis skin. Nature Communications 12 (2021). 10.1038/s41467-021-24607-6

35 Bielli, A. et al. Cellular retinoic acid binding protein-II expression and its potential role in skin aging. Aging 11, 1619–1632 (2019). 10.18632/aging.101813

36 Deng, C.-C. et al. Single-cell RNA-seq reveals immune cell heterogeneity and increased Th17 cells in human fibrotic skin diseases. Frontiers in Immunology 15 (2025). 10.3389/fimmu.2024.1522076

37 Ma, F. et al. Single-cell profiling of prurigo nodularis demonstrates immune-stromal crosstalk driving profibrotic responses and reversal with nemolizumab. Journal of Allergy and Clinical Immunology 153, 146–160 (2024). 10.1016/j.jaci.2023.07.005

38 Walker, J. T., McLeod, K., Kim, S., Conway, S. J. & Hamilton, D. W. Periostin as a multifunctional modulator of the wound healing response. Cell Tissue Res 365, 453–465 (2016). 10.1007/s00441-016-2426-6

39 Yamaguchi, Y. Periostin in Skin Tissue Skin-Related Diseases. Allergol Int 63, 161–170 (2014). 10.2332/allergolint.13-RAI-0685

40 Andersen, M. K. et al. Spatial transcriptomics reveals strong association between SFRP4 and extracellular matrix remodeling in prostate cancer. Communications Biology 7 (2024). 10.1038/s42003-024-07161-x

41 Mauri, F. et al. NR2F2 controls malignant squamous cell carcinoma state by promoting stemness and invasion and repressing differentiation. Nature Cancer 2, 1152–1169 (2021). 10.1038/s43018-021-00287-5

42 Al Harthi, F. et al. Apolipoprotein E Gene Polymorphism and Serum Lipid Profile in Saudi Patients with Psoriasis. Disease Markers 2014, 1–8 (2014). 10.1155/2014/239645

43 Huang, S., Howington, M. B., Dobry, C. J., Evans, C. R. & Leiser, S. F. Flavin-Containing Monooxygenases Are Conserved Regulators of Stress Resistance and Metabolism. Frontiers in Cell and Developmental Biology 9 (2021). 10.3389/fcell.2021.630188

44 Shoffner-Beck, S. K. et al. Lupus dermal fibroblasts are proinflammatory and exhibit a profibrotic phenotype in scarring skin disease. JCI Insight 9 (2024). 10.1172/jci.insight.173437

45 Mizumoto, N. & Takashima, A. CD1a and langerin: acting as more than Langerhans cell markers. 113, 658–660 (2004). 10.1172/jci21140

46 Hilligan, K. L. & Ronchese, F. Antigen presentation by dendritic cells and their instruction of CD4+ T helper cell responses. Cellular & Molecular Immunology 17, 587–599 (2020). 10.1038/s41423-020-0465-0

47 Nierkens, S., Tel, J., Janssen, E. & Adema, G. J. Antigen cross-presentation by dendritic cell subsets: one general or all sergeants? Trends in Immunology 34, 361–370 (2013). 10.1016/j.it.2013.02.007

48 Xia, T. et al. Advances in the study of macrophage polarization in inflammatory immune skin diseases. Journal of Inflammation 20 (2023). 10.1186/s12950-023-00360-z

49 Almlöf, J. C., et al. Novel risk genes for systemic lupus erythematosus predicted by random forest classification. Scientific Reports 7 (2017). 10.1038/s41598-017-06516-1

50 An, X. J. et al. RasGRP3 in peripheral blood mononuclear cells is associated with disease activity and implicated in the development of systemic lupus erythematosus. Am J Transl Res 11, 1800–1809 (2019).

51 Cui, Y., Sheng, Y. & Zhang, X. Genetic susceptibility to SLE: recent progress from GWAS. J Autoimmun 41, 25–33 (2013). 10.1016/j.jaut.2013.01.008

52 He, C. F. et al. TNIP1, SLC15A4, ETS1, RasGRP3 and IKZF1 are associated with clinical features of systemic lupus erythematosus in a Chinese Han population. Lupus 19, 1181–1186 (2010). 10.1177/0961203310367918

53 Blauvelt, A. et al. Next Generation PDE4 Inhibitors that Selectively Target PDE4B/D Subtypes: A Narrative Review. Dermatology and Therapy 13, 3031–3042 (2023). 10.1007/s13555-023-01054-3

54 Crowley, E. L. & Gooderham, M. J. Phosphodiesterase-4 Inhibition in the Management of Psoriasis. Pharmaceutics 16, 23 (2023). 10.3390/pharmaceutics16010023

55 Li, H., Zuo, J. & Tang, W. Phosphodiesterase-4 Inhibitors for the Treatment of Inflammatory Diseases. Frontiers in Pharmacology 9 (2018). 10.3389/fphar.2018.01048

56 Croft, M. et al. OX40 in the Pathogenesis of Atopic Dermatitis—A New Therapeutic Target. American Journal of Clinical Dermatology 25, 447–461 (2024). 10.1007/s40257-023-00838-9

57 Lé, A. M. & Torres, T. OX40-OX40L Inhibition for the Treatment of Atopic Dermatitis—Focus on Rocatinlimab and Amlitelimab. Pharmaceutics 14, 2753 (2022). 10.3390/pharmaceutics14122753

58 Rindler, K. et al. Spontaneously Resolved Atopic Dermatitis Shows Melanocyte and Immune Cell Activation Distinct From Healthy Control Skin. Frontiers in Immunology 12 (2021). 10.3389/fimmu.2021.630892

59 Sadrolashrafi, K. et al. An OX-Tra’Ordinary Tale: The Role of OX40 and OX40L in Atopic Dermatitis. Cells 13, 587 (2024). 10.3390/cells13070587

60 B. Brandt, E. Th2 Cytokines and Atopic Dermatitis. Journal of Clinical & Cellular Immunology 02 (2011). 10.4172/2155-9899.1000110

61 Facheris, P., Jeffery, J., Del Duca, E. & Guttman-Yassky, E. The translational revolution in atopic dermatitis: the paradigm shift from pathogenesis to treatment. Cellular & Molecular Immunology 20, 448–474 (2023). 10.1038/s41423-023-00992-4

62 Ha, H.-L. et al. IL-17 drives psoriatic inflammation via distinct, target cell-specific mechanisms. Proceedings of the National Academy of Sciences 111, E3422–E3431 (2014). 10.1073/pnas.1400513111

63 Mosca, M. et al. The Role of IL-17 Cytokines in Psoriasis. ImmunoTargets and Therapy Volume 10, 409–418 (2021). 10.2147/itt.s240891

64 Pavel, P. et al. Peroxisomal Fatty Acid Oxidation and Glycolysis Are Triggered in Mouse Models of Lesional Atopic Dermatitis. JID Innov 1, 100033 (2021). 10.1016/j.xjidi.2021.100033

65 Yang, W. et al. Meta-analysis Followed by Replication Identifies Loci in or near CDKN1B, TET3, CD80, DRAM1, and ARID5B as Associated with Systemic Lupus Erythematosus in Asians. The American Journal of Human Genetics 92, 41–51 (2013). 10.1016/j.ajhg.2012.11.018

66 Schafer, P. H. et al. Phosphodiesterase 4 in inflammatory diseases: Effects of apremilast in psoriatic blood and in dermal myofibroblasts through the PDE4/CD271 complex. Cell Signal 28, 753–763 (2016). 10.1016/j.cellsig.2016.01.007

67 Marsh, S. scCustomize: Custom Visualizations & Functions for Streamlined Analyses of Single Cell Sequencing., <10.5281/zenodo.5706430> (2021).

68 Hua, Y., Weng, L., Zhao, F. & Rambow, F. SeuratExtend: Streamlining Single-Cell RNA-Seq Analysis Through an Integrated and Intuitive Framework (Cold Spring Harbor Laboratory, 2024).

69 Bunis, D. G., Andrews, J., Fragiadakis, G. K., Burt, T. D. & Sirota, M. dittoSeq: universal user-friendly single-cell and bulk RNA sequencing visualization toolkit. Bioinformatics 36, 5535–5536 (2021). 10.1093/bioinformatics/btaa1011

70 Jin, S., Plikus, M. V. & Nie, Q. CellChat for systematic analysis of cell–cell communication from single-cell transcriptomics. Nature Protocols 20, 180–219 (2025). 10.1038/s41596-024-01045-4

