## Supplementary figures and images for "A Transcriptomic Atlas of Healthy Human Skin Links Regional Identity to Inflammatory Disease"

### Supplementary Figures 1-3

**Supplementary Figure 1**

A.

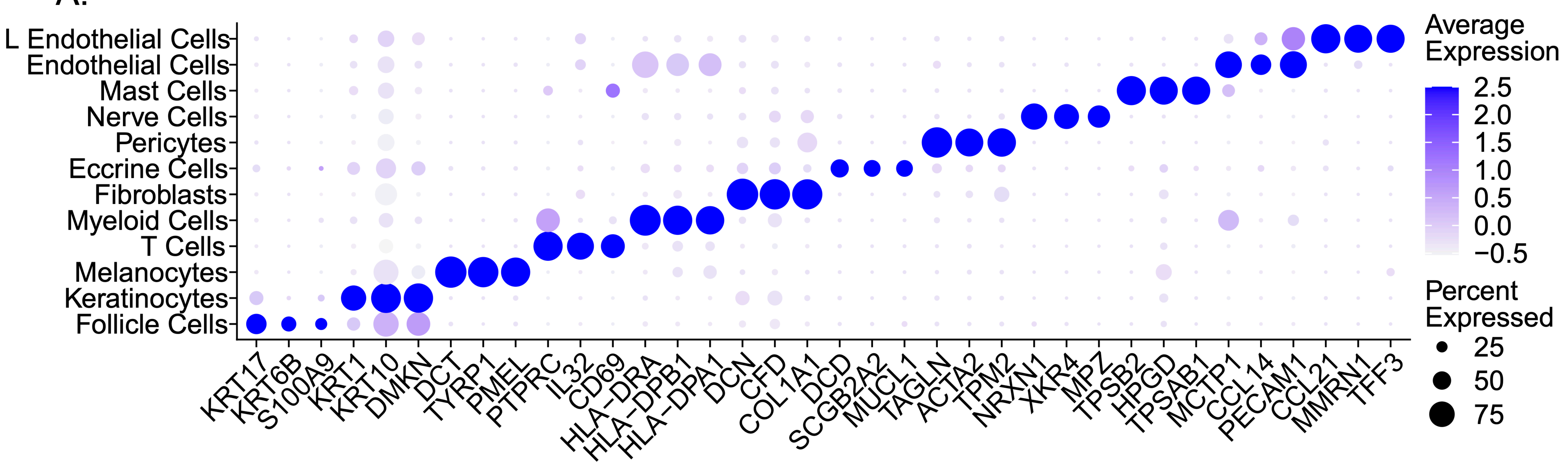


B.

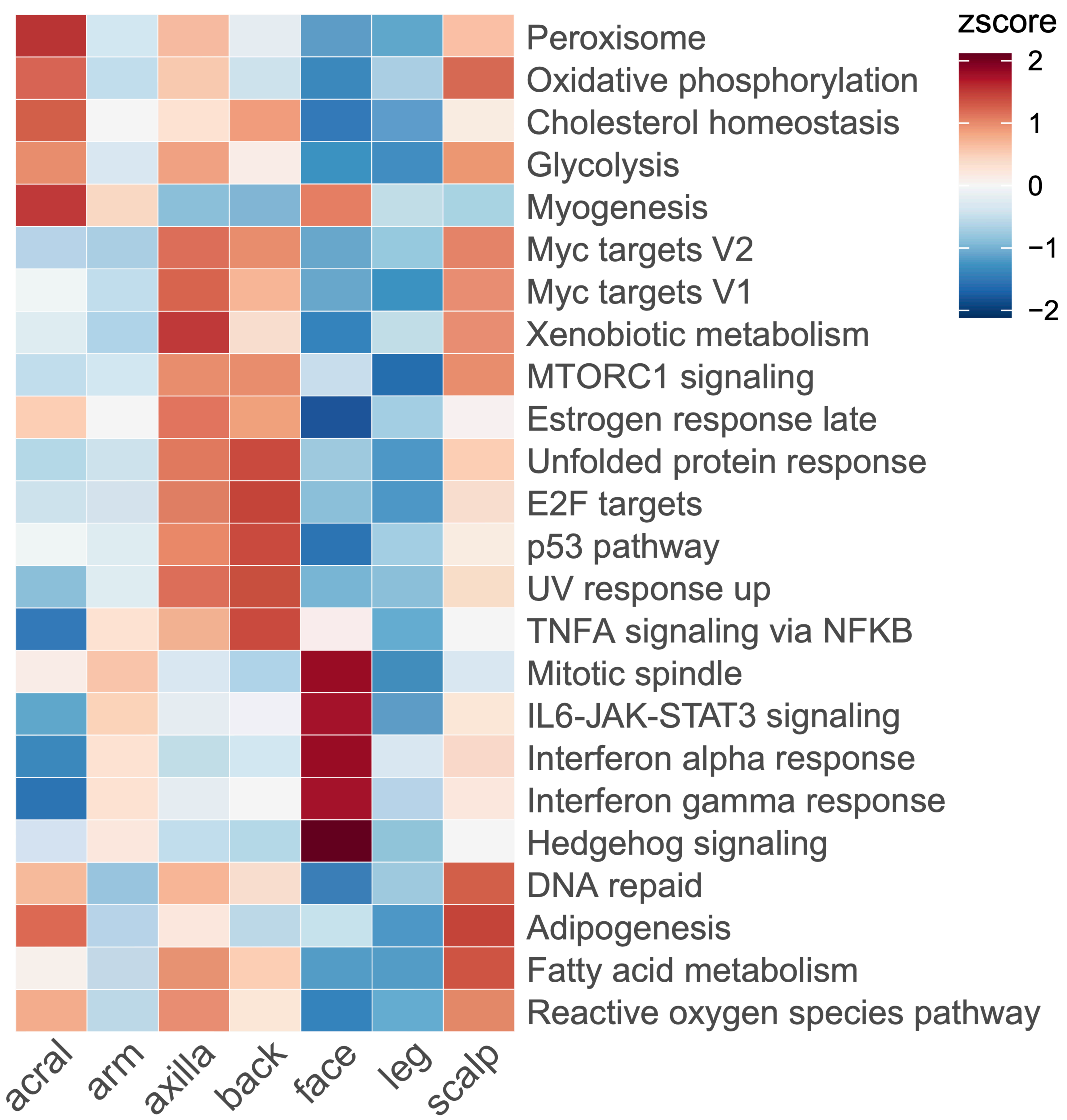

**Supplementary Figure 2**

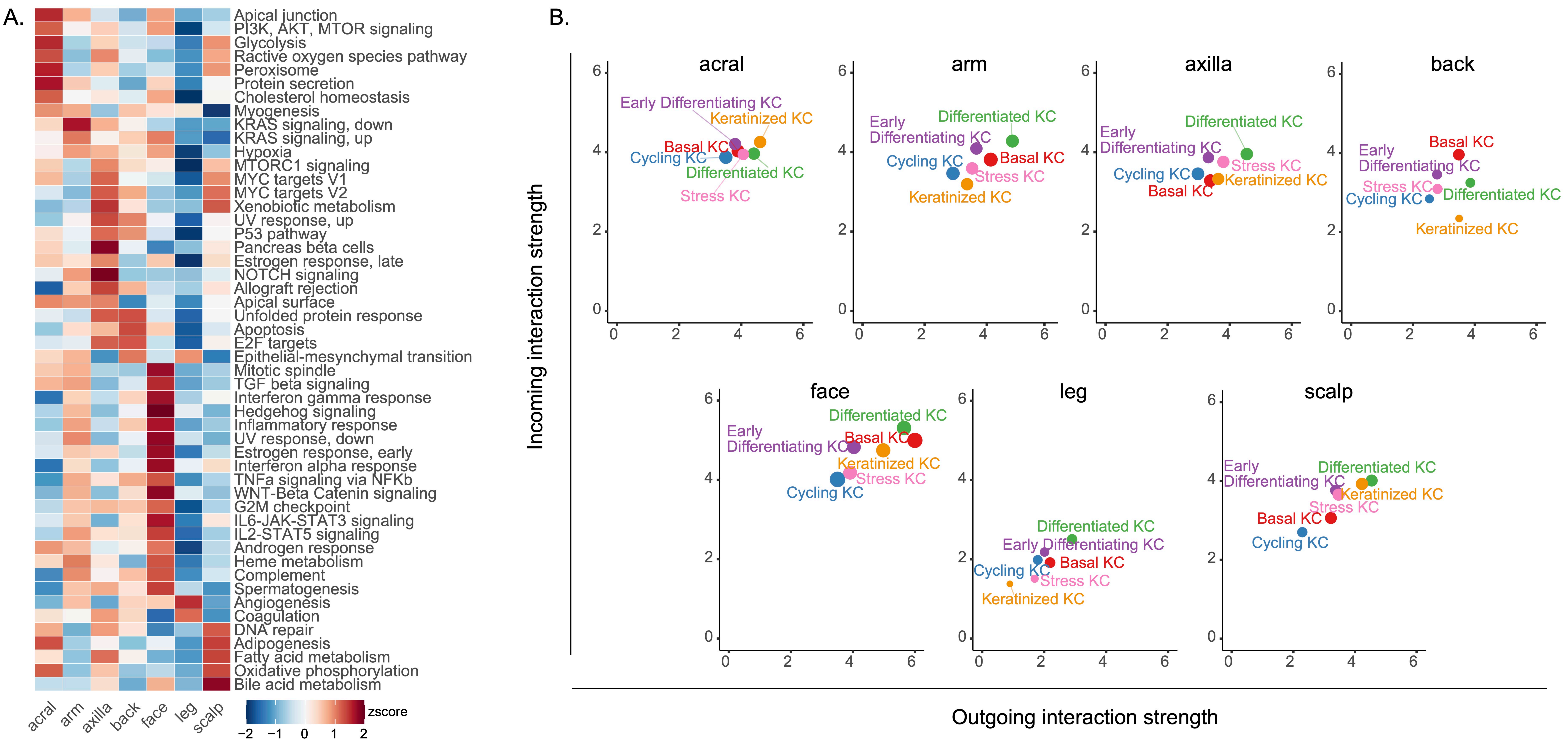

#### Supplementary Figure 3

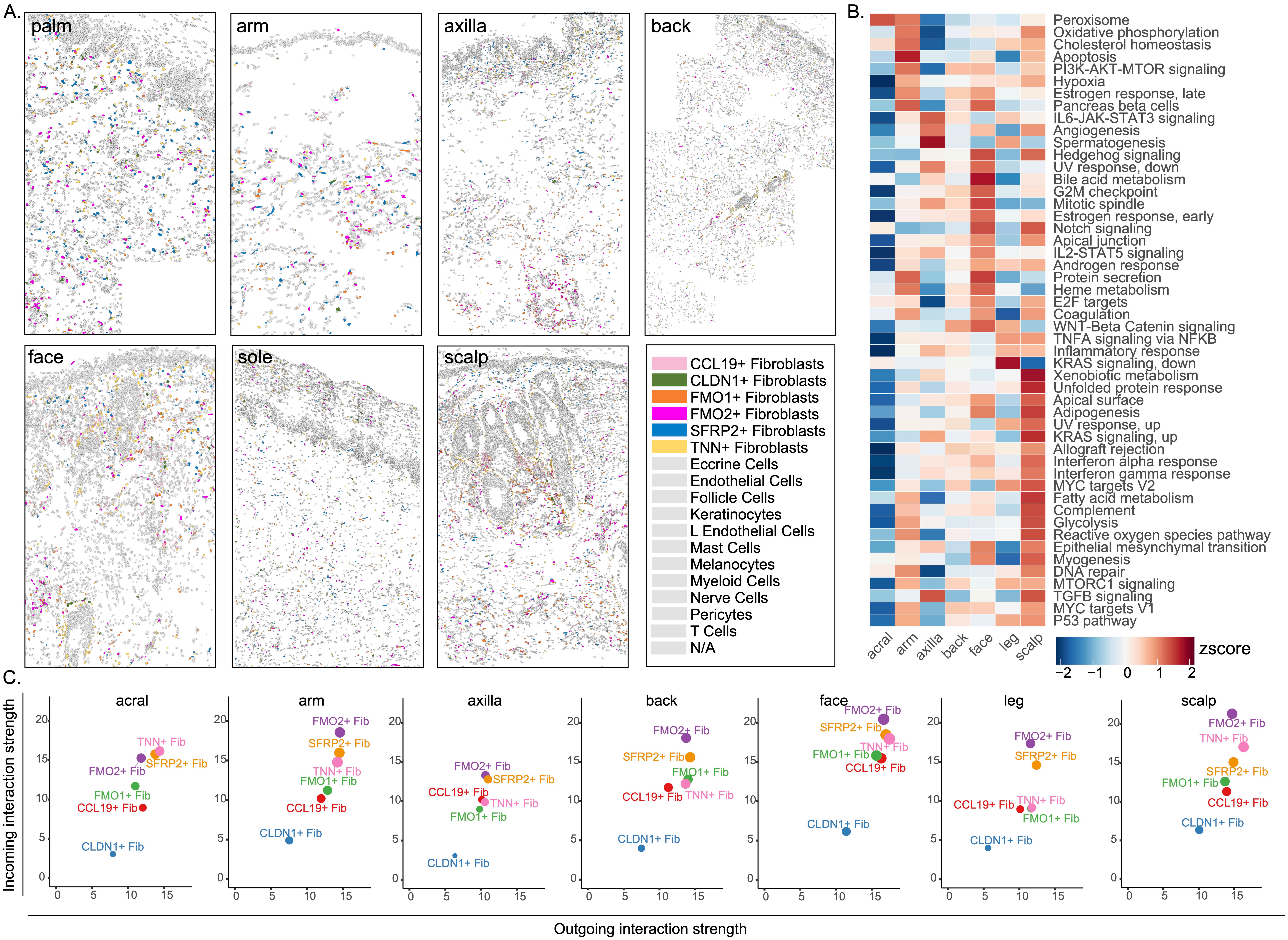
